# Basal epidermis collective migration and local Sonic hedgehog signaling promote skeletal branching morphogenesis in zebrafish fins

**DOI:** 10.1101/2020.06.29.165274

**Authors:** Joshua A. Braunstein, Amy E. Robbins, Scott Stewart, Kryn Stankunas

## Abstract

Teleost fish fins, like all vertebrate limbs, comprise a series of bones laid out in characteristic pattern. Each fin’s distal bony rays typically branch to elaborate skeletal networks providing form and function. Zebrafish caudal fin regeneration studies suggest basal epidermal-expressed Sonic hedgehog (Shh) promotes ray branching by partitioning pools of adjacent pre-osteoblasts. This Shh role is distinct from its famous Zone of Polarizing Activity role establishing paired limb positional information. Therefore, we investigated if and how Shh signaling similarly functions during developmental ray branching of both paired and unpaired fins while resolving cellular dynamics of branching by live imaging. We found *shha* is expressed uniquely by basal epidermal cells overlying pre-osteoblast pools at the distal aspect of outgrowing juvenile fins. Lateral splitting of each *shha*-expressing epidermal domain followed by the pre-osteoblast pools precedes overt ray branching. We use *ptch2:Kaede* fish and Kaede photoconversion to identify short stretches of *shha*+ basal epidermis and juxtaposed pre-osteoblasts as the Shh/Smoothened (Smo) active zone. Basal epidermal distal collective movements continuously replenish each *shha*+ domain with individual cells transiently expressing and responding to Shh. In contrast, pre-osteoblasts maintain Shh/Smo activity until differentiating. Smo inhibition using the small molecule BMS-833923 prevents branching in all fins, paired and unpaired, with surprisingly minimal effects on caudal fin initial skeletal patterning, ray outgrowth or bone differentiation. Staggered BMS-833923 addition indicates Shh/Smo signaling acts throughout the branching process. We use live cell tracking to find Shh/Smo restrains the distal movement of basal epidermal cells by apparent ‘tethering’ to pre-osteoblasts. We propose short-range Shh/Smo signaling promotes these heterotypic associations to couple instructive basal epidermal collective movements to pre-osteoblast repositioning as a unique mode of branching morphogenesis.

## INTRODUCTION

The diversity and beauty of fish fins has long captivated aquarists while providing compelling models to consider morphological evolution and organ patterning. Fins’ intricate skeletons, comprised of elongated dermal bony rays and body-proximal endochondral bones, support swimming biomechanics and define appendage shape. Most rays, or lepidotrichia, branch one or more times to elaborate the skeletal network. Fin rays, as well as ancestral unpaired medial fins, were lost during the fish-to-tetrapod transition (Freitas et al., 2014). Paired pectoral and pelvic fins and their endochondral bones evolved into tetrapod limbs. However, fin rays and limb digits may share deep evolutionary homology (Nakamura et al., 2016). Comparing skeletal patterning between paired and unpaired fins, including the prominent caudal fin, can reveal core appendicular development mechanisms and those underlying evolutionary innovations.

*Danio rerio* zebrafish are widely studied teleost fish with branched rays in all paired and unpaired fins. For example, zebrafish caudal fins typically have 18 rays of which the central 16 branch into “daughter” rays (Figure S1A). Juvenile fish form primary branches around 30 days post fertilization (dpf) followed by secondary and tertiary branches. Individual rays comprise two opposed hemi-rays that form cylindrical skeletal units segmented by joints and enveloped by a multilayered epidermis (Figure S1B). Zebrafish fully regenerate adult fins including restoring a branched ray skeleton within two weeks of injury. Empowered by versatile genetic and other tools, zebrafish have become a leading model for appendage regeneration research. Collective studies implicate many of the same signaling pathways, including Wnt, Bmp, Fgf, retinoic acid, and Hh, involved in tetrapod limb development (Reviewed in Sehring and Weidinger, 2019). Therefore, zebrafish fin development and regeneration provide complementary contexts to understand how cell signaling patterns the appendicular skeleton and how the same pathways are reactivated for repair. Further, zebrafish fins and their rays enable studies of fundamental developmental questions, including how branched networks form – a common property of many organs including vasculature, lungs, kidneys, mammary glands, and pancreas.

Sonic hedgehog (Shh) signaling through its Smoothened (Smo) effector is one pathway dually involved in fin regeneration and appendage development. Shh is associated closely with tetrapod limb skeletal patterning (Zuniga, 2015) and Shh/Smo pathway perturbations cause syndactyly and polydactyly (Anderson et al., 2012; Malik, 2012). Shh is the Zone of Polarizing Activity (ZPA) morphogen that pre-patterns the limb bud into distinct skeletal units, including digits (Chiang et al., 2001; Riddle et al., 1993; Saunders and Gasseling, 1968). Zebrafish *shha* expression studies suggest a ZPA patterns pectoral and other paired fins but not unpaired fins including the caudal fin (Hadzhiev et al., 2007; Laforest et al., 1998). Nevertheless, *shha* is expressed in distal epidermal domains overlying each forming ray during caudal fin development and regeneration (Armstrong et al., 2017; Hadzhiev et al., 2007; Laforest et al., 1998; Lee et al., 2009; Zhang et al., 2012). At least during regeneration, each *shha*-expressing domain splits prior to ray branching. We leveraged the highly specific Smo inhibitor BMS-833923 (BMS) to show Shh/Smo specifically promotes ray branching during zebrafish caudal fin regeneration (Armstrong et al., 2017). However, a potentially similar Shh/Smo role during developmental ray branching of all fins and underlying mechanisms are unresolved.

During caudal fin regeneration, distal-moving basal epidermal cells (bEps) adjacent to bony rays upregulate *shha* at the distal “progenitor zone” (Armstrong et al., 2017; Figure S1C). Shh-expressing bEps activate Hedgehog/Smoothened (Hh/Smo) signaling in themselves and immediately adjacent pre-osteoblasts (pObs) as marked by *patched2* (*ptch2*), which encodes a Hh receptor and universal negative feedback regulator (Alexandre et al., 1996; Goodrich et al., 1996; Lorberbaum et al., 2016; Marigo et al., 1996). This short-range Hh/Smo signaling is required to split pOb pools and therefore ray branching without impacting proliferation or differentiation. While Hh/Smo signaling is active continuously, *shha*-expressing basal epidermal domains split laterally prior to pOb partitioning (Figure S1D). We proposed Shh/Smo signaling might enhance physical associations between moving epidermal cells and pObs to enable the progressive partition of pOb pools (Armstrong et al., 2017). However, this model has not been assessed during fin development or expanded on by visualization of cell movement dynamics in live animals.

Here, we explore the mechanisms underlying developmental fin ray branching in both paired and unpaired fins of juvenile zebrafish. We show basal epidermal dynamics as well as Shh/Smo activity and function are largely the same as during regeneration. We use transgenic reporter lines for *shha* and its target gene *ptch2* to refine developmental expression profiles. Kaede photoconversion of *TgBAC(ptch2:Kaede)* fish reveals continuous Shh/Smo signaling in distal ray Shh+ basal epidermal domains and neighboring pObs. We inhibit Shh/Smo signaling using the small molecule BMS-833923 to show the pathway is largely dedicated to ray branching in all fins, including paired fins. Shh+ bEps and pObs are closely apposed at the site of Shh/Smo signaling where a basement membrane is incompletely assembled. bEps constantly move distally, trafficking through while contributing to *shha*-expressing domains that split laterally prior to ray branching. We use live time-lapse imaging to demonstrate Shh/Smo signaling restrains basal epidermal distal migration, possibly by transiently coupling *shha*+ bEps to distal ray pObs. We conclude the collective migration of bEps, constantly distal with progressive lateral domain splitting, and their atypical use of local Shh/Smo signaling re-positions pObs for skeletal branching during both fin development and regeneration. This reflects a unique branching morphogenesis process whereby movements of a neighboring cell type – the bEps – guides the tissue-forming cells – the pObs – into split pools.

## RESULTS

### Shha expression is progressively restricted to distal ray basal epidermal domains that split preceding ray branching

Developing fins express *sonic hedgehog a* (*shha*) in single basal epidermal domains adjacent to the tip of each hemi-ray up to 20 days post fertilization (dpf) (Hadzhiev et al., 2007; Laforest et al., 1998). During caudal fin regeneration, similar *shha*-positive basal epidermal domains split into distinct domains around 4 days post amputation (dpa) immediately preceding ray branching (Armstrong et al., 2017; Laforest et al., 1998; Lee et al., 2009; Quint et al., 2002; Zhang et al., 2012). We expected a comparable pattern during fin development if branching during fin regeneration recapitulates developmental processes. We used the reporter line *Tg(-2.4shha:gfpABC)^sb15^,* which mimics endogenous *shha* expression (*shha:GFP*; (Ertzer et al., 2007; Zhang et al., 2012), to monitor *shha* expression in live zebrafish from its emergence through primary ray branching in juveniles around 30 dpf. *shha:GFP* expression in the caudal region was restricted to the developing floor plate and notochord through 5 dpf, as previously established (Krauss et al., 1993). From 5-9 dpf, *shha:GFP* expression expanded into the caudal fin fold primordium through a ventral gap of melanophores (Figure 1A, white arrowheads). By 9-10 dpf, *shha:GFP*+ cells were enriched along emerging immature rays (Figure 1B). At 10-12 dpf, *shha:GFP* expression became specifically associated with the distal aspect of maturing central rays while remaining along the lengths of immature peripheral rays (Figure 1C). *shha:GFP* was restricted to ray tips by 14 dpa, when all rays contained maturing, segmented bone (Figure 1D).

**Figure 1.**
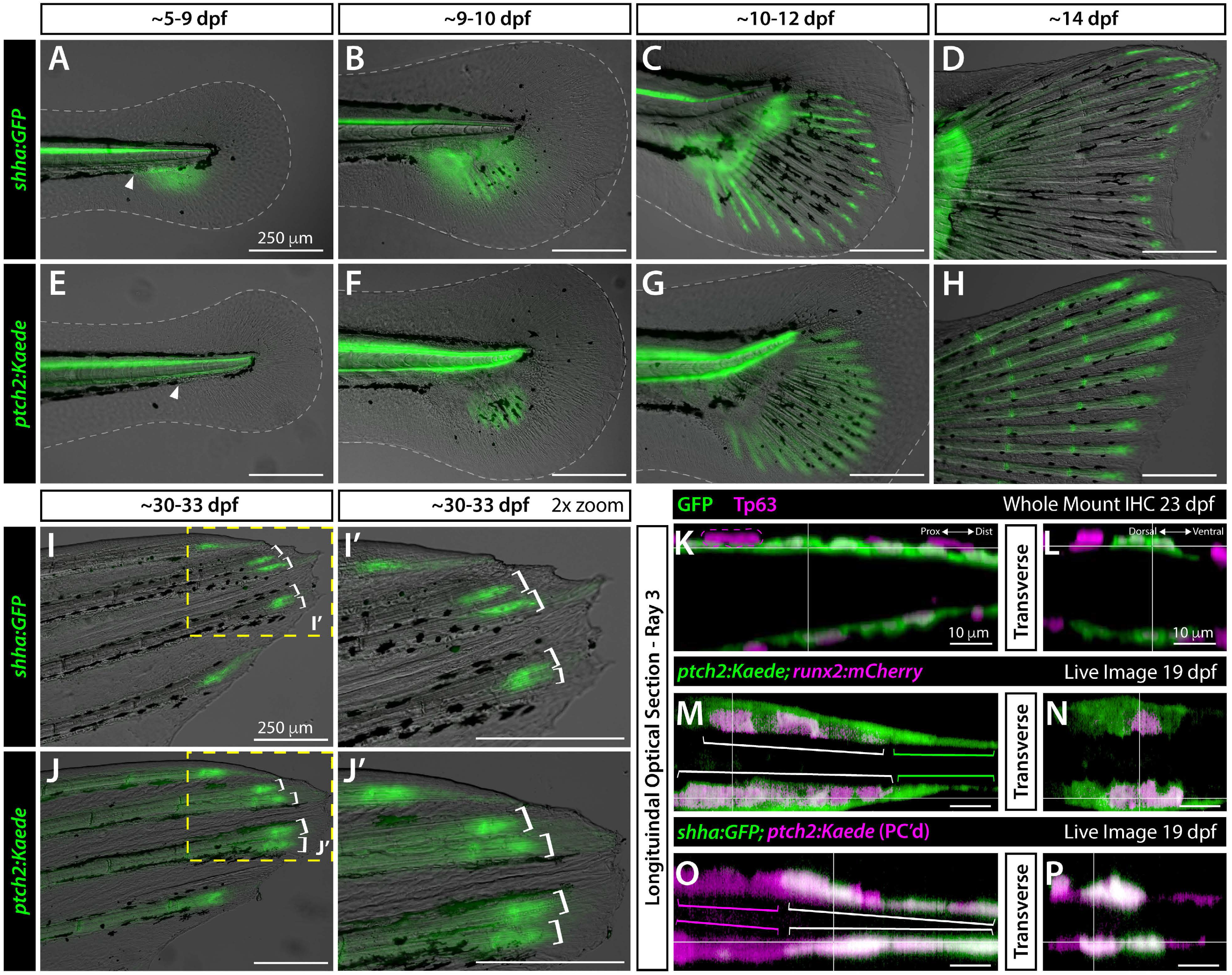
Basal epidermal *shha* and *ptch2*-defined responses in basal epidermis and pre-osteoblasts become progressively distally restricted during caudal fin development. (A-J’) Differential interference contrast and fluorescent overlay images of developing caudal fins of *shha:GFP* (A-D, I, I’) and *ptch2:Kaede* (E-H, J, J’) fish of indicated ages. White dotted lines outline the fin fold (A-F). The white arrowhea indicates the gap in melanophores where *shha:GFP* expression emerges. Yellow boxes in (I, J) indicate the 2x zoom fields in (I’, J’). White brackets (I-J’) mark branched reporter domains in dorsal rays 2 and 3 preceding overt ray branching. (K-P) Single optical slices of caudal fin dorsal ray 3 in longitudinal (K, M, O) and transverse (L, N, P) planes derived from 3D-reconstructed whole mount confocal images of fluorescent reporter fish of indicated ages. (K, L) 23 dpf *shha:GFP* fin whole mount antibody stained for GFP (green) and the basal epidermis marker Tp63 (magenta). The magenta dashed line outlines a representative Tp63+, GFP-cell that occasionally overlay the innermost basal epidermal layer (magenta dashed outline, K). (M-P) Single optical slice reconstructed equivalents from live whole mount-imaged 19 dpf *ptch2:Kaede;runx2:mCherry* and *shha:GFP;ptch2:Kaede* caudal fins. (M, N) *ptch2:Kaede* (green) is in distal *runx2*-marked pre-osteoblasts (magenta; white brackets) and a thin layer of tightly associated adjacent and further distally-extending basal epidermis (green brackets). (O, P) *ptch2:Kaede* (photoconverted; magenta) in pre-osteoblasts (magenta brackets) and co-localized with *shha:GFP*-expressing basal epidermis (green, white brackets). Scale bars are 250 μm in (A-J’) and 10 μm in (K-P).

The *shha:GFP* domains began splitting around 30 dpf, immediately prior to branching of the corresponding ray (Figure 1I, I’, white brackets). We used whole mount immunostaining for GFP and the basal epidermal marker Tp63 (Lee and Kimelman, 2002) and 3D confocal reconstructions to confirm *shha*-driven GFP was expressed exclusively in basal epidermal cells (bEps) (Figure 1K, L; expanded data in Figure S2). Where Tp63-marked cells were multi-layered (magenta dashed line, Figure 1K), only the innermost cells expressed GFP. Similar *shha* expression patterns, including *shha*+ bEp domain splitting, support a common Shh/Smo-dependent mechanism for both developmental and regenerative ray branching.

Zebrafish have 3 unpaired (dorsal, anal, and caudal) and 4 paired (pectoral and pelvic) fin appendages, each with a branched dermoskeleton. We explored if *shha:GFP* domains split ahead of developmental ray branching in all fins. We imaged dissected fins from 51 dpf juvenile fish carrying *shha:GFP* and *Tg(runx2:mCherry)* (*runx2:mCherry*; Shannon Fisher Lab, unpublished; Barske et al., 2020), which marks *runx2+* pre-osteoblasts (pObs) and then perdures in differentiated Obs along ray lengths. *shha:GFP* was expressed at and slightly distal to the *runx2:mCherry+* pOb*-*defined fin boundary in all seven fins, including paired pectoral and pelvic fins (Figure S3). In all cases, branching rays were tipped with two distinct *shha:GFP+* domains, suggesting shared Shh/Smo-promoted ray branching regardless of fin evolutionary or morphological divergence.

### Ptch2 expression indicates Hh/Smo activity in basal epidermis and pre-osteoblasts

We next assessed expression of *ptch2*, a Hh/Smo negative feedback regulator and activity marker (Alexandre et al., 1996; Goodrich et al., 1996; Lorberbaum et al., 2016; Marigo et al., 1996). Hh/Smo signaling induces *ptch2* in pObs and neighboring bEps during caudal fin regeneration (Armstrong et al., 2017; Quint et al., 2002). *ptch2* is also expressed in distal regions of late larval caudal fins, although cell-level expression is unresolved (Laforest et al., 1998). We imaged the *TgBAC(ptch2:kaede)^a4596^* reporter line (*ptch2:Kaede*; Huang et al., 2012) at the same developmental time points we examined *shha:GFP*. *ptch2:Kaede* expression was confined to the notochord and floor plate until ∼9 dpf (Figure 1E). By 9-10 dpf, *ptch2:Kaede* was associated with nascent rays in the ventrally expanding fin primordial (Figure 1F). At 10-12 dpf, *ptch2:Kaede* was detected the entire length of each ray (Figure 1G). This pattern, which includes past *ptch2* expression due to prolonged Kaede stability, persisted through 14 dpf with notably higher *ptch2:Kaede* expression associated with joints and distal ray tips, matching its pattern in regenerating fins (Figure 1H; Armstrong et al., 2017). As with *shha:GFP*, *ptch2:Kaede+* domains split as ray branching initiated in 30-33 dpf juvenile fish (Figure 1J, J’).

To define the cell types expressing *ptch2* during caudal fin development, we combined *ptch2:Kaede* with *runx2:mCherry* and *shha:GFP* to mark *runx2*+ pObs and *shha*+ bEps, respectively (Figure 1M-P and expanded data with single channel images in Figure S4). Confocal imaging of live 19 dpf double transgenic larval fish showed *ptch2:Kaede* co-localized with distal fin *runx2:mCherry*-expressing pObs (Figure 1M, N). Only adjacent and further distal presumptive bEps additionally expressed *ptch2:Kaede.* We used photoconversion to discern Kaede from GFP and reveal non-pOb *ptch2:Kaede* co-localized with *shha:GFP*-expressing and nearby bEps (Figure 1O, P). Therefore, *ptch2* defines autocrine (in bEps) and short-range (in pObs and non-*shha*+ bEps) Shh/Smo signaling during caudal fin development.

### Active Shha/Smo signaling is restricted to outgrowing distal ray regions

We photoconverted distal ray ends of 25 dpf *ptch2:Kaede* caudal fins and re-imaged 24 hours later to distinguish actively produced Kaede from perduring reporter fluorescence (Figure 2A-C). New, unconverted Kaede was produced exclusively in short, discrete domains at the ray tips. Therefore, active Shh/Smo signaling appears narrowly focused in close proximity with *shha*-expressing bEps. We also observed a stretch of photoconverted Kaede+ cells distal to the ray tips in tissue newly formed over the 24-hour post-conversion period. We saw a similar pattern during fin regeneration and likewise identify the cells as distal-moving bEps given their broader domain and more distal location than ray-forming osteoblasts (Figure S5F’; Armstrong et al., 2017). Therefore, previously Shh-responsive bEps rapidly cease *ptch2* expression as they collectively move beyond the Shh/Smo active zone.

**Figure 2.**
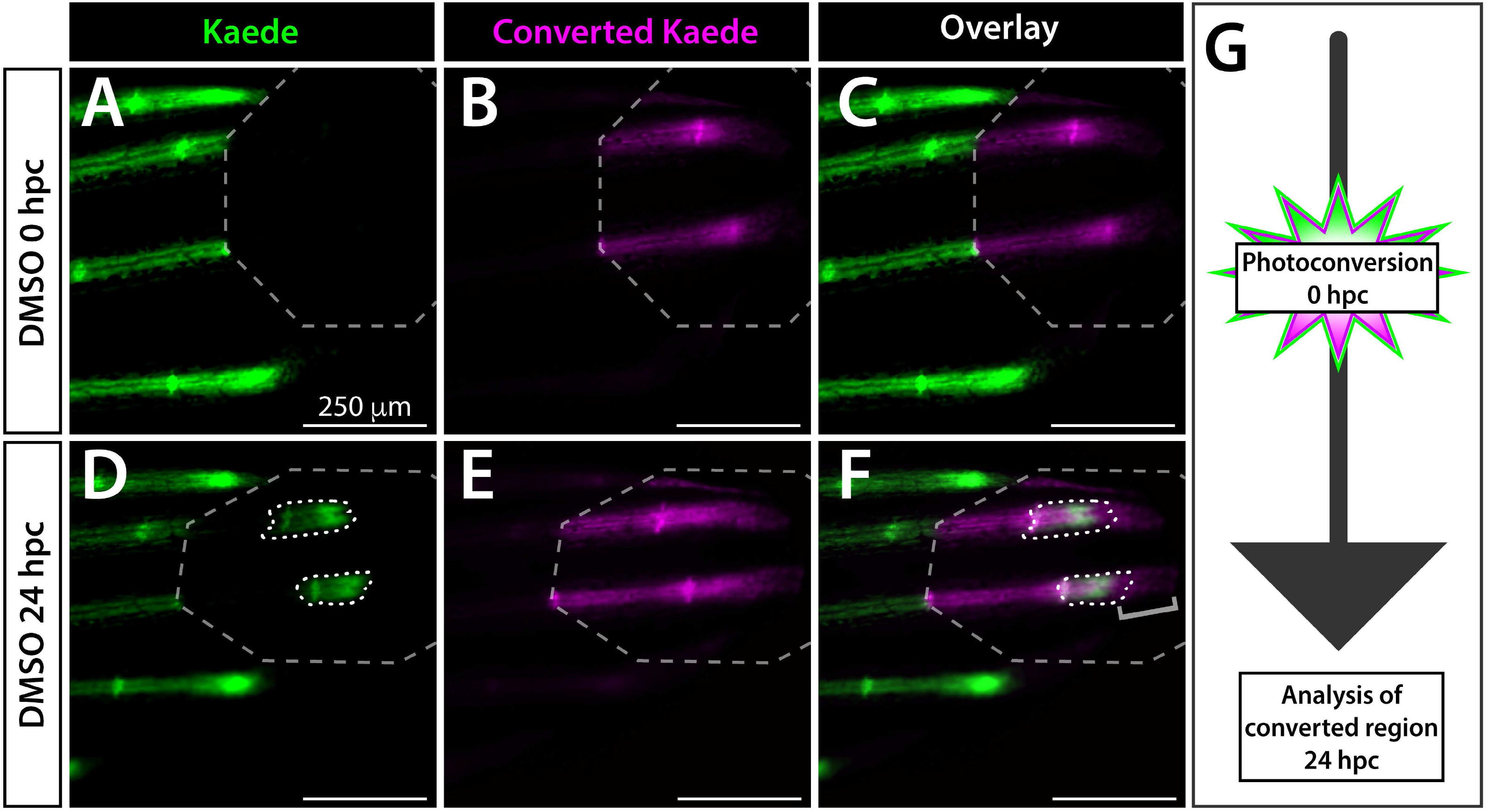
Active Shh/Smo signaling is restricted to a narrow distal stretch of each developing fin ray. (A-F) Whole mount fluorescent images of the distal aspect of the caudal fin of a 25 dpf *ptch2:Kaede* larval fish at the time of Kaede photoconversion from green to red (false colored magenta) fluorescence (0 hpc, A-C) and 24 hours later (24 hpc, D-F). The grey dashed octagon marks the photoconverted region of interest (ROI). The 0 hpc overlay (C) demonstrates complete Kaede photoconversion. The same fish imaged at 24 hpc displays a small patch of newly produced green Kaede (white dotted outlines in D, F) within the photoconverted ROI. The grey bracket in (F) marks distally displaced basal epidermis retaining photoconverted Kaede. Slight splitting of the new green Kaede domain (D, F) indicates the onset of ray branching. (G) Schematic of the photoconversion time course. The fish shown is representative of *n=*8. Scale bars are 250 μm.

We used the potent Smo inhibitor BMS-833923 (Armstrong et al., 2017; Lin and Matsui, 2012; henceforth abbreviated BMS) to confirm *ptch2:Kaede* reports Shh/Smo activity during fin development. As expected, caudal fins of 25 dpf fish treated with 1.25 µM BMS and photoconverted 3 hours later produced no new Kaede 24 hours post-conversion (hpc; Figure S5G-L). Curiously, we no longer observed distally displaced photoconverted Kaede+ bEps with the remaining photoconverted Kaede narrowly associated with ray tips and therefore likely pObs (Figure S5F’, L’). We surmise the 24-hour Shh/Smo-inhibition led distal-moving Kaede+ bEps to shed prematurely, leaving only photoconverted Kaede+ pObs. As such, Shh/Smo signaling may retard distal bEp collective movements.

### Sustained Shh/Smo signaling promotes ray branching during fin development

We next investigated if Shh/Smo is required for ray branching during fin development. Treating *Tg(sp7:EGFP)^b1212^* osteoblast reporter fish (*sp7:EGFP*; DeLaurier et al., 2010) with 0.63 µM BMS from 25 to 42 dpf blocked caudal fin ray branching (Figure 3A-D, *n*=5 per BMS and DMSO control groups). In contrast, Shh/Smo-inhibition did not disrupt caudal fin outgrowth or skeletal maturation, indicated by well-defined cylindrical bony ray segments complete with joints, of the central 16 rays (Figure S6). Curiously, the non-branching principal peripheral rays uniquely were shorter in BMS-treated fish (Figure 3A, C white arrows and Figure S6). BMS-treatment of 29 dpf *shha:GFP* fish exposed to EdU for 12 hours did not change the fraction of EdU+ intra-ray cells, i.e. pObs and mesenchyme nestled between the epidermal Shh domains of each hemi-ray (Figure S7, *n*=5 per group). We conclude Shh/Smo signaling is largely dedicated to ray branching with minimal proliferation or bone maturation effects during both fin development and, as shown previously, regeneration.

**Figure 3.**
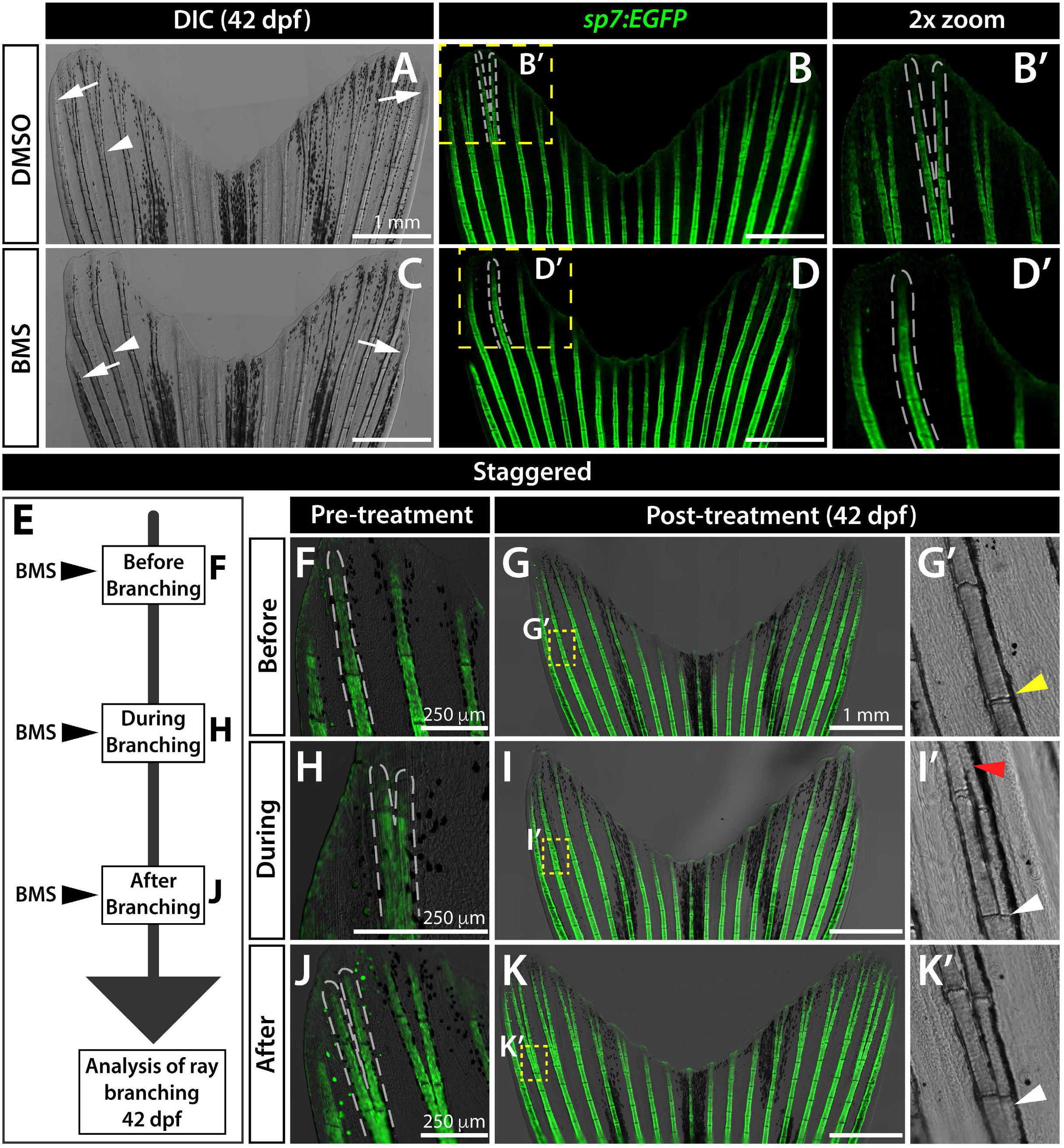
Sustained Shh/Smo signaling is required for ray branching in developing caudal fins. (A-D) Differential interference contrast (DIC) and fluorescence images of caudal fins from DMSO (A, B) and BMS-833923 (BMS)-treated (C, D) 42 dpf *sp7:EGFP* osteoblast reporter fish. Dashed white lines outline dorsal ray 3 and daughter rays, when present. White arrowheads designate a present (A) or absent (B) branch point. White arrows mark ends of the principal peripheral rays. (E-K) Experimental schematic of (E) and resulting caudal fin images of 42 dpf *sp7:EGFP* fish from (F-K’) staggering the start of BMS treatment to before (F-G’), during (H-I’), and after branch initiation (J-K’). Yellow dashed boxes outline dorsal ray 2 regions shown in (G’, I’ and K’). The yellow arrowhead in (G’) designates where branching would have occurred without Shh/Smo-inhibition. The red arrowhead in (I’) marks where a ray re-fused when BMS was added after branching had initiated. White arrowheads in (I’ and K’) indicate ray branch points. Images represent treatment groups (before, during, and after) of *n=*4 or 5 BMS-treated and *n* =2 or 3 DMSO controls. Ray re-fusions occur in 4 of 5 fish treated with BMS “during” ray branching in rays 2, 3, and 16. Scale bars are 250 μm or 1 mm, as indicated.

Shh/Smo signaling may act transiently to initiate ray branching or continuously during the branching process. To distinguish between these possibilities, we staggered the start of BMS treatment to “before”, “during”, or “after” branching (Figure 3E), identified by *a priori* screening 24-35 dpf *sp7:EGFP* clutchmate fish. Expectedly, BMS-exposure initiated prior to ray branching prevented said rays from branching (“before” group, Figure 3F, G, G’, *n*=4/4) and rays that had already fully branched remained so after BMS treatment (“after” group, J, K, K’, *n*=4/4). However, rays that recently initiated branching (“during” group, Figure 3H) re-fused upon BMS exposure, forming “gapped” ray segments (Figure 3I, I’, *n*=4/5). Therefore, sustained Shh signaling acts throughout ray branching morphogenesis rather than as a switch that initiates branching.

### Shh/Smo signaling does not substantially contribute to initial fin ray patterning

*shha* and *ptch2* expression during early stages of caudal fin formation (Figure 1B, E), while non-polarized, is reminiscent of Shh’s ZPA role in paired appendages. This pattern suggests Shh/Smo may influence early caudal fin skeletal patterning in addition to promoting later, juvenile-stage ray branching. To explore this possibility, we inhibited Shh/Smo signaling from as early as 2 dpf, when the larval fin fold entirely comprises soft tissue absent of any ray structures. As expected, *ptch2:Kaede*-marked Shh/Smo signaling was restricted to the notochord and floor plate at this stage (Figure S8A-C). Photoconversion experiments confirmed BMS fully inhibited production of new *ptch2:Kaede* in 14 dpf larval caudal fins (Figure S8D-E’, total *n*=33-44 per group), as with embryos, juvenile fins, and regenerating adult fins (Figure S4G-L; (Armstrong et al., 2017). We treated *sp7:EGFP;runx2:mCherry* fish with 1.25 μM BMS from 2 until 14 dpf, when all 18 rays were clearly established. Their caudal fins developed the standard complement of 18 rays with a central diastema and apparently normal length distribution across the dorsal-ventral axis to define typical “V-shaped” fins (Figure S8F-G’). As such, Shh/Smo may have limited or no role in pre-patterning the caudal fin field despite early *shha* and *ptch2* expression.

### Shh/Smo signaling supports ray branching in all fins

We tested if Shh/Smo signaling promotes ray branching in all seven fins, paired and unpaired, by treating *shha:GFP;runx2:mCherry* fish with BMS starting at 21 dpf, prior to asynchronous ray branching across fins. As expected, both DMSO control and BMS-treated fish showed *shha:GFP+* domains at the distal end of every ray of all fins at 42 dpf (Figure 4). However, BMS-treated fish showed no or, at best, severely delayed branching in all fins (*n*=6 per group). The rare, delayed branching likely reflects incomplete Shh/Smo inhibition due to the challenge of sustaining effective drug concentrations across a multi-week treatment. Nevertheless, the near absence of branchpoints confirms all fins employ a common Shh/Smo signaling-dependent mechanism for ray branching irrespective of their evolutionary divergence, including the apparent *shha*-defining ZPA in paired fins (Hadzhiev et al., 2007; Laforest et al., 1998).

**Figure 4.**
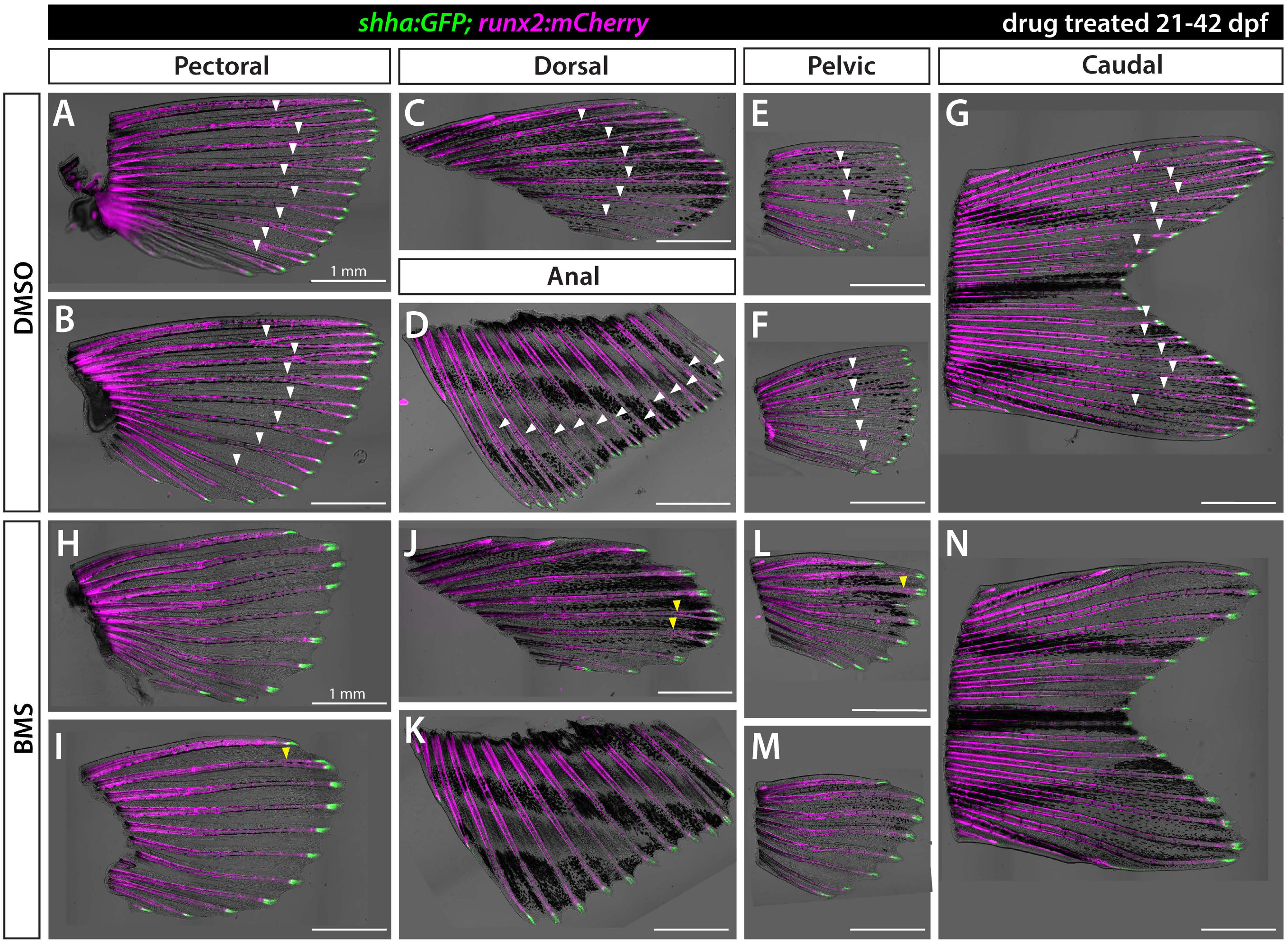
Shh/Smo signaling promotes ray branching in all paired and unpaired fins. Brightfield and fluorescence overlay images of isolated fins from *shha:GFP;runx2:mCherry* juvenile fish treated with DMSO (A-G) or 1.25 µM BMS-822923 (BMS; H-N) from 21-42 dpf. *runx2:mCherry*-labeled rays branch (white arrowheads) in all fins whereas rays of BMS-treated fish mostly fail to branch or have severely delayed branching (yellow arrowheads). *shha:GFP+* basal epidermal cells are restricted to distal ray tips under both conditions. *n=*6 for each group. Scale bars are 1 mm.

### Shha*+* bEps and pObs are intimately associated in developing caudal fins

We next aimed to identify how sustained, local Shh/Smo signaling affects bEp and/or pOb cell behaviors to promote ray branching morphogenesis. The close proximity of these two Shh-responsive cell types suggested their movements might be physically coupled in a Shh/Smo-dependent manner to promote branching. To assess potential physical contacts between bEps and pObs, we first stained longitudinal sections of 32 dpf juvenile fin rays from *shha:GFP* fish with GFP, the osteoblast marker Zns-5, and Laminin, a component of the epidermal-osteoblast separating basement membrane. As expected, Shha:GFP+ bEps were directly adjacent to pObs (Figure S9). A thin Laminin-containing basement membrane separated pObs and the proximal-most Shha:GFP+ bEps that had recently arrived in the active zone and initiated *shha* expression. More distally, the double staining for Shha:GFP+ bEps and Zns-5+ pObs produced even partially overlapping signal (Figure S9D and D’), suggesting the two cell types are intimately associated. Here, the Laminin+ basement membrane was less dense and sometimes fragmented, likely reflecting its nascent production (asterisks, D’).

We further explored the relative positioning of bEps and pObs at the onset of ray branching by 3D confocal reconstructions of live imaged fins of 28 dpf *shha:GFP;runx2:mCherry* fish (Figure 5A-C). *shha:GFP+* bEps and *runx2:mCherry+* pObs were tightly juxtaposed in both hemi-rays of a single lepidotrichia (Movie 1). Focusing on one hemi-ray, we observed extensive apparent heterotypic contacts, including areas where *shha:GFP+* bEps enshrouded a ridge of pObs (Movie 2). Single sagittal optical slices and reconstructed slice equivalents examined multi-dimensionally (Figure 5D-F) showed intertwined bEps and pObs unresolvable by conventional confocal microscopy.

**Figure 5.**
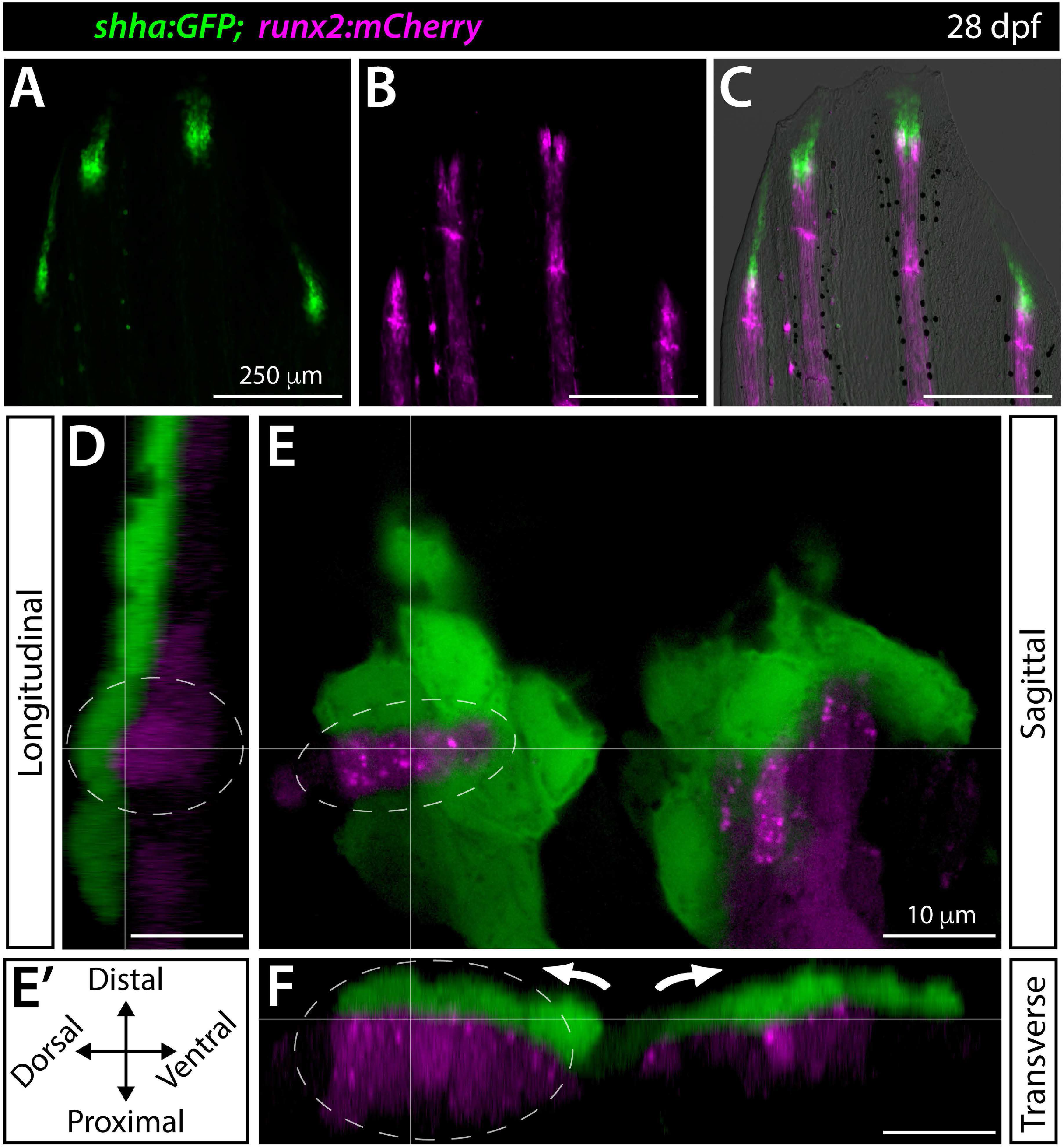
Shha+ basal epidermis and pre-osteoblasts are intertwined in developing fins. (A-F) Fluorescence widefield (A-C) or confocal (D-F) images of the dorsal caudal fin lobe of a 28 dpf *shha:GFP;runx2:mCherry* fish. (A-C) *shha:GFP*-expressing basal epidermal cells (green) overlay and extend distally from *runx:mCherry*-high pre-osteoblast (magenta). The overlay in (C) includes a brightfield image for context. (D-F) A single optical slice (E; sagittal; orientation key in E’) and reconstructed longitudinal (D) and transverse (F) views of a distal ray region undergoing ray branching. *shha:GFP+* basal epidermis (green) and *runx2:mCherry+* pre-osteoblasts (magenta) have overlapping signal at interfaces, indicating their tight juxtaposition. Basal epidermis and pre-osteoblasts tandemly separate into split pools during branching (white arrows). The grey dotted oval highlights a ridge of pre-osteoblasts nestled into a *shha:GFP+* basal epidermal groove (Movie 2). Scale bars are 250 μm (A-C) and 10 μm (D-F).

We considered if Shh/Smo signaling promotes the juxtaposition of bEps and pObs. However, BMS treatment of *shha:GFP;runx2:mCherry* fish from 24-34 dpf did not alter the intimate association between Shh+ bEps and Runx2+ pObs in static images of live fins even though the same drug exposure prevented ray branching to 42 dpf (Figure S10). As expected, Runx2+ pOb pools failed to split upon BMS exposure, even when *shha:GFP* domains still did. The *shha:GFP* domains of BMS-treated fish variably appeared to remain as one cluster per hemi-ray (Figure S10E, J; 4/10 split for dorsal ray 3). In contrast, our fin regeneration study indicated Shha-domain splitting is always Shh/Smo-independent (Armstrong et al., 2017). The difference could reflect difficulty visualizing the initial splitting of the smaller *shha:GFP* domains of juvenile fins. Further, constant *shha* production throughout fin growth, not just at branching, seemingly precludes a direct Shh/Smo role in periodically splitting bEp domains. Regardless, Shh/Smo signaling does not support ray branching by promoting close proximity between bEps and pObs per se.

### Shh/Smo signaling restrains basal epidermal collective movements while adjacent to pObs

Shh/Smo inhibition appeared to increase the rate of bEp shedding due to accelerated distal collective movements (Figure S5I, L). Therefore, we considered if Shh/Smo transiently couples bEps and pObs by direct cell-to-cell adhesion or through intermediary connections that impede bEp movements while they neighbor relatively stable pObs. Such regulated heterotypic associations, which may not be evident by static imaging, coupled with force-generating bEp collective movements during *shha*+ domain splitting, could re-position pOb pools over time. We time-lapse imaged caudal fins of *ptch2:Kaede* fish at late larval stages (22-24 dpf) to assess heterotypic cellular dynamics in outgrowing rays. *ptch2:Kaede+* bEps moved distally over *ptch2:Kaede+* pObs, which remained stationary over the 30 minute imaging period, and in more distal fin tissue (Movie 3, Figure S11A-C). We used semi-automated tracking of individual *ptch2:Kaede+* bEps to determine *ptch2:Kaede+* bEps of BMS treated fish moved significantly faster (3-6 cells per fish and *n*=8 fish per group) (Figure S11D). Therefore, Shh/Smo signaling restrains the distal movement of Shh/Smo-responsive bEps.

We imaged caudal fins of *shha:GFP*;*runx2:mCherry* larval fish with and without Shh/Smo inhibition to monitor movement dynamics of *shha:GFP*-expressing bEps relative to Runx2+ pObs (representative fish in Figure 6A-D’ and Movie 4; all fish shown in Figure S12). We resolved individual cells at higher detail in distal ray regions by capturing full confocal z-stacks every 2 minutes over 30 minutes. Assisted by semi-automated cell tracking, we noted bEps moved faster when beyond the field of pObs in DMSO-treated control fins. We observed slow moving bEps in contact with pObs before detaching and rapidly moving distally. In contrast, BMS exposure caused rapid distal bEp movement irrespective of proximal-to-distal position or proximity to pObs.

**Figure 6.**
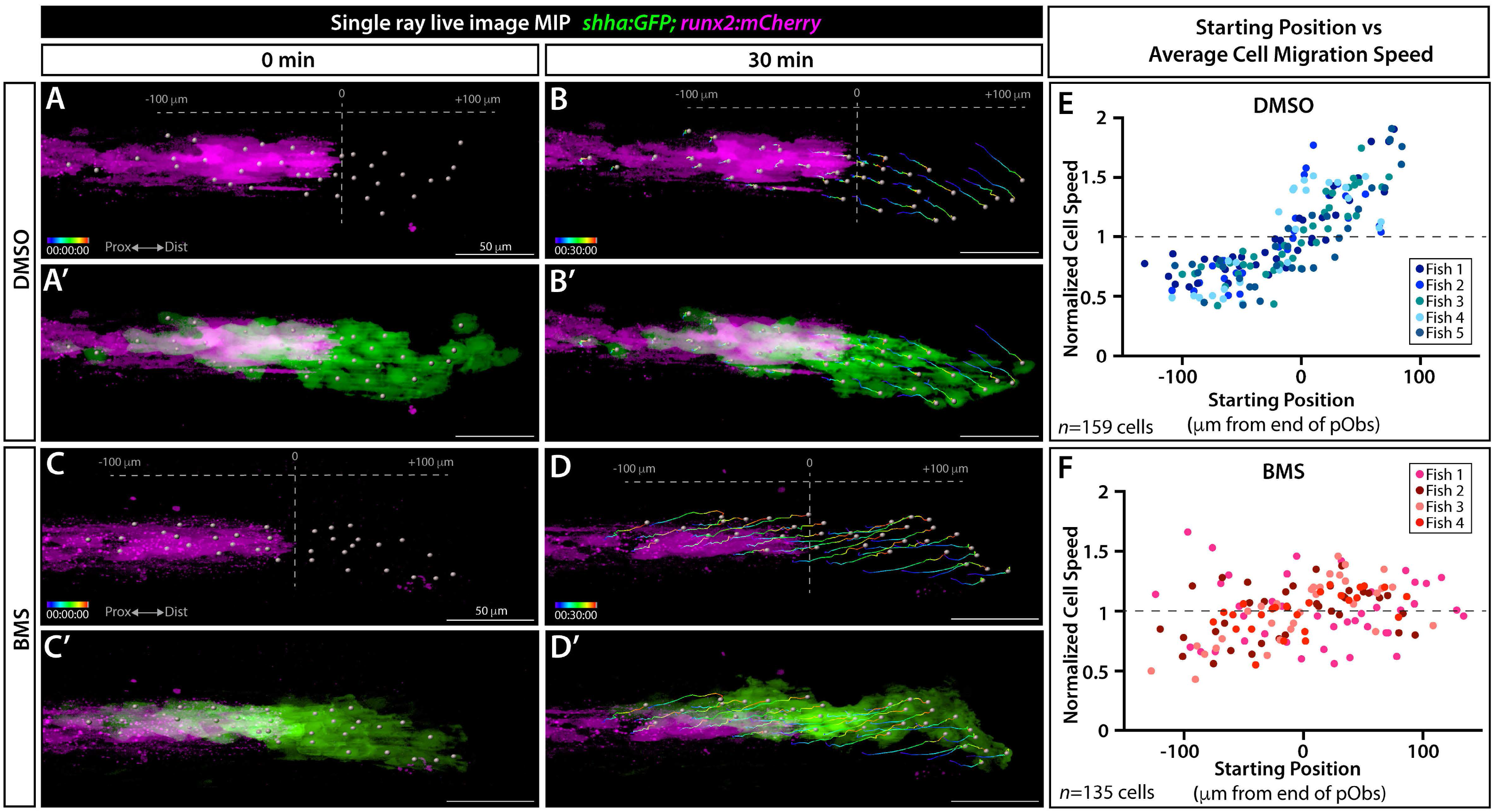
Shh/Smo signaling slows collective migration of *shha*-expressing basal epidermal cells associated with pre-osteoblasts. (A-D’) Frames from a time lapse movie showing dorsal ray 3 of the caudal fin from live mounted 24 dpf *shha:GFP;runx2:mCherry* fish treated with DMSO or BMS-833923 (BMS) for 24 hours prior to imaging. Images are maximum intensity projections (MIP) and show the start (0 min; A, A’, C, C’) and end (30 min; B, B’, D, D’) points. The Imaris-generated colored tracks show the progressive displacement of individual *shha:GFP+* basal epidermal cells (green). Grey spheres show the starting or final positions of all tracked basal epidermal cells. Grey dashed vertical lines in (A, B, C, D) indicate the distal most Runx2+ pre-osteoblast (magenta), defined as position “0”. (E, F) Scatterplot graphs showing the average speed of individual basal epidermal cells, considering their net X-displacement and normalized to all scored cells of the given fish, relative to starting position for DMSO-(159 individual cells from five fish) and BMS-exposed fish (135 cells from four fish). Dot colors correspond to cells from a given fish. Scale bars are 50 μm.

We quantified positional dynamics of individual bEps and plotted their average normalized speed compared to starting position relative to the end of ray-forming Runx2+ pObs (DMSO: *n*=5 fish, 26-38 cells per fish, total of 159 cells; BMS: *n*=4 fish, 26-41 cells per fish, total of 135 cells). *shha:GFP*-expressing bEps located distal to pObs moved faster than pOb-associated bEps in control animals, producing a clear upward velocity shift at the pOb border (Figure 6E). In contrast, BMS treatment caused evenly distributed bEp velocities before and after the pOb-containing region. Taken together, we propose local Shh/Smo signaling enhances heterotypic cell associations that transiently restrain the continuous distal movement of bEps while they pass directly over pObs. For ray branching morphogenesis, Shh/Smo-enhanced cell interactions between pObs and successive waves of bEps could enable the pOb pool to gradually follow laterally splitting *shha*-expressing bEp domains. Eventually, the divided pOb pools would then form separate daughter rays connected at a branch point.

## DISCUSSION

### Basal epidermal movements and Shh/Smo signaling direct skeletal branching morphogenesis during zebrafish fin ray development and regeneration

Zebrafish fin ray branching provides an accessible context to define ancestral mechanisms of appendicular skeletal morphogenesis. Our current and earlier study (Armstrong et al., 2017) extends previous research to demonstrate the same Shh-dependent branching morphogenesis mechanism branches developing and regenerating rays. In both contexts, a gradually splitting domain of Shh-expressing basal epidermal cells (bEps) at the distal aspect of each outgrowing fin ray partitions the immediately adjacent pre-osteoblast (pOb) pool. Highly localized, continuous Shh/Smo activity allows a given ray’s distal pOb population to gradually follow the separating Shha+ basal epidermal domains. Eventually fully split, divided pOb pools continue to promote outgrowth of now two rays connected at a branch point. The shared mechanism of pOb positioning for ray branching underscores that fin regeneration re-activates developmental mechanisms. We show Shh/Smo-promoted ray branching morphogenesis acts in unpaired and paired fins, distinct from the paired fins’ presumptive earlier use of *shha* in a ZPA-like patterning role (Hadzhiev et al., 2007; Laforest et al., 1998). Smo-dependent Shh signaling appears largely dedicated to ray branching in unpaired fins as pathway inhibition minimally disrupts initial fin patterning, outgrowth, or skeletal differentiation during caudal fin development or regeneration.

Live imaging of the developing caudal fin highlights how collective movement of Shha+ bEps positions pObs to generate branched rays (model in Figure S13 and Movie 5). Our Kaede photoconversion and time-lapse imaging show bEps continuously move distally in developing fins, activating *shha* expression upon reaching the distal zone that includes pObs. Individual bEps pass through the *shha*-expressing domain, down-regulate *shha* when moving beyond the pObs, and then are seemingly shed from the end of the fin. In turn, proximal bEps enter the distal zone and activate *shha* to replenish the *shha*-expressing basal epidermal domain adjacent to pObs. Shha produces a constant Smo-dependent response in neighboring pObs and an autocrine, transient response in *shha*-producing bEps as represented by upregulated *ptch2* in both cell types. This continuous, localized Hh/Smo signaling restrains bEp collective movement dynamics and promotes ray branching by enabling concomitant separation of pOb pools with lateral splitting *shha* basal epidermal domains.

The directly observed distal movement and likely shedding of fins’ basal epidermis, as surmised during regeneration (Armstrong et al., 2017; Shibata et al., 2018) is intriguing both functionally and mechanistically. Functionally, continuous bEp replacement may enable rapid recovery from frequent environmental insults. Further, we propose bEp collective movements, distal and lateral, contribute the force that maintains pOb alignment (Armstrong et al., 2017) and enables splitting of physically coupled pOb pools for ray branching. Distal epidermal movements are likely promoted by distributed proliferation across the fin, as seen during regeneration for the basal epidermis (Shibata et al., 2018) and overlying superficial epidermis (Chen et al., 2016; Shibata et al., 2018). In contrast, the cause of periodic lateral *shha*-expressing bEp movements that split *shha*-defined bEp domains and thereby instruct ray branching is unknown.

Shh/Smo signaling is involved in branching morphogenesis of other organs, including the lung (Bellusci et al., 1997; Fernandes-Silva et al., 2017; Pepicelli et al., 1998) and submandibular salivary gland (Jaskoll et al., 2004). In the lung, Shh/Smo mediates interactions between mesenchymal and epithelial populations although likely by promoting local proliferation and/or differentiation (Kim et al., 2015). Collective cell migration also is broadly implicated in branching morphogenesis, including for renal tubes, mammary glands and blood vessels (Ewald et al., 2008; Riccio et al., 2016; Spurlin and Nelson, 2017). Unlike those contexts, we propose a neighboring cell type – basal epidermis – that does not directly contribute to the final tissue provides the instructive collective movements. This unusual arrangement may reflect regenerating fin pObs having a mesenchymal state during patterning before returning to their differentiated epithelial state (Stewart et al., 2014).

### Shh/Smo signaling may position pre-osteoblasts by physical coupling to moving basal epidermal cells

Continuous Shh/Smo signaling through fin development and retained splitting of Shh-expressing basal epidermal domains when the pathway is inhibited indicate Shh/Smo has a permissive role in ray branching. We favor a model whereby Shh/Smo’s function is to promote transient associations between bEps and pObs (Figure S13). The transient nature may result from the moving bEps rapidly terminating their *shha* expression and *ptch2*-defined Shh/Smo activity. A slight lateral component to bEp movements away from the midline of each forming ray would then successively tug interconnected pObs to follow. Over the course of several days, pObs eventually are pulled into two pools. The pOb pools become sufficiently and irreversibly separated to now generate branched daughter rays.

Our 3-D reconstructions showing Shh-expressing bEps and Runx2-expressing pObs likely share extensive and intimate physical contacts are consistent with this heterotypic cell association model. Importantly, bEps and pObs would remain adjacent but not necessarily physically coupled when Shh/Smo signaling is blocked. As such, our novel time-lapse imaging of developing caudal fins provides key functional support by showing Shh/Smo signaling impedes bEp distal movements (Movie 5). Notably, Shh-expressing bEps accelerate when they pass beyond Shh-responding pObs. Chemical inhibition of Shh/Smo signaling significantly increases overall Ptch2-positive bEp migration rates and eliminates the characteristic velocity decrease when Shh-expressing bEps pass adjacent to pObs. While inhibiting Shh/Smo signaling accelerates individual bEp cell movements, a steady-state Shh-expressing basal epidermal domain persists and at least partially splits. However, we propose the pObs cannot follow without Shh/Smo signaling to couple them with bEps and therefore remain as a single pOb pool that forms an unbranched ray. Longer-term live imaging monitoring lateral movements of both bEps and pObs would help test this model.

We favor the model Shh/Smo physically couples bEps and pObs movements for ray branching over alternative hypotheses for additional reasons. First, we did not observe Shh/Smo-dependent cell proliferation during development or regeneration (Armstrong et al., 2017) arguing against a model whereby Shh promotes localized proliferation at the margins of a given pOb pool to progressively divide it. Notably, previous conclusions that Shh/Smo signaling is a pro-proliferative factor in regenerating fins (Lee et al., 2009; Quint et al., 2002) may reflect off-target cyclopamine effects (Armstrong et al., 2017). In contrast, fins develop and regenerate to normal size when using BMS-833923 to block Shh/Smo signaling with the intriguing exception of the two principal peripheral rays. Second, all Shh/Smo-responsive cells remain outwardly specified upon Shh/Smo inhibition, including osteoblasts that still differentiate to produce ray skeletal units complete with joints. Any additional Shha or other Hedgehog ligand roles appear minor or may be Smo-independent, as with Indian Hedgehog A (Ihha) and bone maturation during fin regeneration (Armstrong et al., 2017). Third, we use chemical genetics to map the Shh/Smo time-of-function and show ray branching requires persistent Shh/Smo signaling from initial hints of Shh-expressing basal epidermal domain splitting until daughter rays are fully separated. Therefore, Shh/Smo signaling seemingly promotes a continuously emergent process and not a periodic switch-like event for either distal bEp movement or Shha-expressing bEp domain splitting.

### Short range Shh promoting cell associations may be a common Shh/Smo signaling mode

Our proposed short range Shh/Smo signaling mode promoting heterotypic cell association differs from Hh’s more typical role as a gradient-forming morphogen. Providing precedence, Hh acts on neighboring cells in several well-established contexts. For example, Hh famously mediates interactions between directly adjacent cells during *Drosophila* embryo segment boundary formation (Ingham, 1993). Short-range Shh/Smo signaling also occurs in vertebrates, including mammalian hair follicle development (Sato et al., 1999; Woo et al., 2012), avian limb patterning (Sanders et al., 2013), and zebrafish retina development (Shkumatava, 2004). Perhaps most germane, *shha+* epidermal cells organize directly underlying Hh-responsive dermal cells during zebrafish scale morphogenesis (Aman et al., 2018).

Shh/Smo has also been tied to cell associations in other settings. For example, Hh’s archetypal role in *Drosophila* wing disc compartment boundary establishment (Ayers et al., 2010) may be through increased “cell bonding” (Rudolf et al., 2015). Shh alters neural crest cell adhesion and migration during avian neural tube morphogenesis (Fournier-Thibault et al., 2009; Jarov et al., 2003; Testaz et al., 2001). Further, misregulated Shh/Smo signaling is linked to invasive cell migration associated with liver, breast, ovarian, and skin cancers (Chen et al., 2013, 2014; Hanna and Shevde, 2016; Zeng et al., 2017).

Identification, characterization, and manipulation of Shh/Smo-upregulated molecules effecting bEp and pOb interactions would strengthen our ray branching model. Such Shh/Smo targets could directly promote cell adhesion, as for neural tube morphogenesis (Jarov et al., 2003; Tsai et al., 2020), or indirectly as extracellular matrix intermediaries. Alternatively, Shh/Smo activity could alter cell features (e.g. shape, polarity, or interconnectivity) that indirectly favor heterotypic association. A third intriguing possibility is that Shh/Smo-upregulated Patched directly binds to membrane-retained Shh on bEp surfaces to increase high-affinity contacts between pObs and bEps. This mechanism could apply elsewhere given Patched is an evolutionary-conserved Shh/Smo-target gene (Alexandre et al., 1996; Goodrich et al., 1996; Lorberbaum et al., 2016; Marigo et al., 1996) while placing Patched in the curious position as both a Shh/Smo effector and negative feedback regulator.

### Fin ray branching as an ancestral mechanism of Shh-mediated appendage patterning and skeletal morphogenesis

How vertebrates pattern skeletal appendages (fins, limbs) is a textbook question of evolutionary and developmental biology. Rays were lost in tetrapod lineages although fin dermal skeleton and tetrapod digits may share deep homology (Nakamura et al., 2016). If so, our demonstration Shh/Smo signaling supports developmental ray branching morphogenesis is intriguing given Shh’s long-appreciated but mechanistically distinct role in vertebrate digit patterning. Shh is the secreted morphogen produced by the zone of polarizing activity (ZPA) at the posterior edge of developing limb buds that directs anterior-to-posterior patterning of skeletal elements including digits (Tickle and Towers, 2017), Polarized *shha* expression in zebrafish pectoral fin buds indicates paired fins employ ZPA-like skeletal patterning (Akimenko and Ekker, 1995; Krauss et al., 1993; Neumann et al., 1999). In contrast, caudal fin primordia lack polarized *shha* (Hadzhiev et al., 2007; Laforest et al., 1998; our results). Consistently, we found disrupting Shh/Smo signaling even prior to formation of the caudal fin field does not alter the initial complement of 18 rays. Moreover, we demonstrate Shh/Smo signaling is required for ray branching in all fins, whether paired or unpaired. The unpaired medial fins (dorsal, caudal, anal) evolved prior to paired fin appendages (Dahn et al., 2006; Desvignes et al., 2018; Freitas et al., 2006; Larouche et al., 2017). Therefore, Shh-dependent ray branching may reflect an ancestral skeletal morphogenesis mechanism that predates emergence of ZPA-based appendage patterning. The waning fitness demand for a ray branching mechanism during the water-to-land transition then may have released selective pressure on the Shh/ZPA module to facilitate terrestrial limb adaptations. Classic models postulating a branching component to endochondral limb skeleton patterning (Oster et al., 1988; Shubin and Alberch, 1986) further inspire exploring if aspects of Shh/Smo’s function during ray branching endure in ZPA-dependent limb patterning.

Shh/Smo signaling promotes skeletal morphogenesis in many contexts beyond limbs. In zebrafish, Shh/Smo patterns craniofacial dermal bones, as illustrated by the opercle (Huycke et al., 2012), and both developing and regenerating scales (Aman et al., 2018). Shh/Smo signaling also impacts mesenchymal cell movements to pattern bird feathers, another albeit non-ossified skin appendage (Li et al., 2018). Shh further supports patterning of the axial skeleton (Chiang et al., 1996; Choi et al., 2012; Dworkin et al., 2016; Hu et al., 2015; Hu and Helms, 1999; Jeong et al., 2004; Swartz et al., 2012) as well as teeth (Ahn et al., 2010; Dassule et al., 2000; Seppala et al., 2017). Our discovery Shh/Smo signaling enables neighboring cells to position pObs during fin ray branching suggests similar mechanisms act in other skeletal patterning contexts. If so, manipulating Shh/Smo pathway to position therapeutically delivered or endogenous progenitor cells could enhance skeletal regenerative medicine.

## MATERIALS AND METHODS

### Zebrafish

*Danio rerio* zebrafish were maintained in 28-29°C circulating fish water within the University of Oregon Aquatic Animal Care Services (UO AqACS) fish facility. Adult zebrafish were housed in Techniplast polycarbonate containers and fed twice daily with dry pellets (“Zebrafish Juvenile Diet”, #388765-134-684, Zeigler Bros., Inc.). Standard housing densities were *n*=25/3.5-liter tank for fish aged > 21 dpf, *n*=25/1.1-liter tank for larvae 4-21 dpf, and *n*=25/100 mL petri dish for < 4 dpf larvae. The following lines were used: wildtype AB, *Tg(sp7:EGFP)^b1212^* (DeLaurier et al., 2010)*, TgBAC(ptch2:Kaede)^a4596^* (Huang et al., 2012), *Tg(-2.4shha:gfp:ABC)^sb15^* [previously known as *Tg(-2.2shh:gfp:ABC)*] (Ertzer et al., 2007; Shkumatava, 2004), (*Tg(runx2:mCherry)* (From Shannon Fisher Lab; Barske et al., 2020). The University of Oregon Institutional Animal Care and Use Committee (IACUC) approved zebrafish experiments.

Fish were staged by days post fertilization (dpf) and visually screened to identify stages of ray branching morphogenesis development (i.e. pre-branching, semi-branched, branched). Approximate standard lengths (SL) for the developmental stages used are listed below:

| Approx. SL (mm) | Approx. dpf | Developmental stage | Caudal fin state |
| --- | --- | --- | --- |
| 3 | 2 | Larval | Fin fold |
| 5-6 | 14 | Mid-larval | Unbranched rays |
| 9-11 | 24-28 | Late larval | Unbranched rays |
| 11.5-13.5 | 28-33 | Juvenile | Active ray branching |
| 14-15 | 35 | Juvenile | Branched rays |
| 20 | 42 | Juvenile | Branched rays |

### Microscopy

Larval and juvenile fish were anesthetized with 74 µg/ml tricaine (MS-222, Syndel) in fish facility system water. Fish or dissected fins were transferred immediately to a 35 mm glass bottom FluoroDish plate (World Precision Instruments). Two or three drops of 1% low-melt agarose, stored at 38°C and cooled before application, were placed on the caudal fin. Fins were quickly flattened to the FluoroDish with a single-hair paintbrush before the agarose hardened. The following microscopes were used: Nikon Eclipse T*i-*E widefield and Nikon Eclipse T*i2-*E with Yokogawa CSU-W1 spinning disk attachments, and Zeiss LSM 880 laser scanning confocal microscope. Confocal image stacks were processed using Imaris software to generate single optical slice digital sections, surface renderings, and 3D reconstructions. Adobe Photoshop was used to adjust levels with identical image acquisition and processing settings for a given experiment. Live fish promptly were euthanized or returned to tanks after imaging.

### Kaede photoconversion and imaging

*ptch2:Kaede* fish were anesthetized and placed on FluoroDish plates as described above. Fins were viewed with a Nikon Eclipse T*i-*E widefield microscope or Nikon T*i*2*-*E/ Yokogawa CSU-W1 spinning disk confocal microscope. Kaede-expressing regions of interest (ROIs) were photoconverted using a metal halide light source and DAPI excitation filter or with 405 nm laser illumination from 10 seconds to 2 minutes, depending on ROI size and fish age. Before and after images were acquired to ensure complete photoconversion of Kaede from green (518 nm) to red (580 nm) emission. Fish were returned to system water and then similarly re-imaged after defined periods. *ptch2:Kaede* expression in different cell types was discerned by fin cells’ distinctive relative positions and morphologies confirmed by co-marker staining (Figure 1M-P and Figure S4).

### BMS-833923 treatments

BMS-833923 (“BMS”, Cayman Chemicals) was dissolved in DMSO to a concentration of 50 mM and single-use aliquots of 6.3-12.5 mM were prepared from stock for each experiment. Aliquots were diluted to a final concentration of 0.63-1.25 µM in system fish water for both larval and juvenile zebrafish treatments. Equal volumes of DMSO were used for control group treatments. Concentrations varied because of batch-dependent drug potency. Each batch of BMS was validated and optimized using photoconversion experiments on *ptch2:Kaede* fish to define a drug concentration that fully inhibited production of Hh/Smo activity-marking new Kaede.

To test Hh/Smo requirements for caudal fin ray branching, 25 dpf *sp7:EGFP* fish (*n*=6 per group) with unbranched rays were treated initially for 24 hours in BMS or DMSO-alone water and then returned to standard housing. Fish were exposed to BMS for 4 hours every other day until the experimental end point at 42 dpf. To assess Hh/Smo roles in all fin appendages, *shha:GFP;runx2:mCherry* fish were treated with 1.25 µM BMS or DMSO (*n*= 6 per group) from 21 to 42 dpf. Fins were dissected and imaged as described above.

For staggered-start juvenile fish treatments, 25 dpf *sp7:EGFP* fish were anesthetized, fluorescently screened, and sorted into groups of those having caudal fins with “unbranched” rays or fins in which branching had initiated but was incomplete (“during”). Fish from the two groups were then treated with BMS as described above. Untreated clutchmate *sp7:EGFP* fish were returned to standard housing and screened every other day until all fish had developed branched rays. Drug treatment of the “branched” group of fish was started at 35 dpf. All treatments ended at 42 dpf, when fins were mounted and imaged as described. For BMS-treated fish (unbranched, during branching, and branched), *n*=4 or 5 fish per group with *n*=2 or 3 for DMSO-treated control groups.

For early larval development studies, *sp7:EGFP;runx2:mCherry* and *ptch2:Kaede* fish were bathed in 1.25 µM BMS starting at 2 dpf alongside DMSO-treated controls (*n*=33-44 fish per group, per clutch). The same drug exposure regiment described for juvenile fish was used. From 2-4 dpf, larvae were treated in 40 mL embryo media in petri dishes. From 5-14 dpf, fish were drug-exposed in beakers containing 125 mL embryo media. Drug efficacy on larval fish was assessed by photoconverting distal fin ROIs of *ptch2:Kaede* fins (photoconversion methods described above) at 13 dpf and re-imaging those regions at 14 dpf (*n*=3-5 per group). All fish were screened for skeletal patterning phenotypes by widefield microscopy. Across clutches, 35/44 (79.5%) BMS-treated larvae developed normally (9/44 or 20.5% were runted) compared to 26/33 (78.8%) DMSO-treated larvae (7/33 or 21.2% were runted). The ∼20% incidence of developmentally delayed larvae was likely caused by extended periods in 250 mL beakers instead of larger nursery tanks. Regardless, nearly all larvae irrespective of size in both groups developed the normal complement of 18 caudal fin rays.

### Ray morphometrics

Ray lengths were assessed for *sp7:EGFP* clutch mates treated from 25-42 dpf with BMS-833923 or DMSO (experiment described above, *n*=6 per group). Using Fiji-ImageJ software, the Principal Peripheral Ray (unbranching lateral ray) and Dorsal Ray 3 (longest branching ray) were measured from 42 dpf endpoint caudal fin images from the proximal base of the fin to the distal fin end. Fin widths were used to normal for body size but did not differ between DMSO and BMS groups. Raw and normalized data were graphed with GraphPad Prism V8 and significance assessed with a Student’s unpaired t-test.

### Whole mount immunostaining

*shha:GFP* caudal fins were harvested at 22-23 dpf and immediately fixed in 4% PFA/PBS overnight at 4°C or for 4 hours at room temperature. Fins were washed extensively in PBS + 0.1% Tween-20 and blocked in 1x PBS, 1% Triton X-100, 5% Normal Goat Serum, and 10% DMSO buffer overnight at 4°C. Fins were incubated with primary antibodies in blocking buffer overnight at 4°C. Primary antibodies were anti-GFP (1:1000; AVES, GFP-1020), anti-Tp63 (1:100; Thermo Fisher, PA5-36069) and anti-Runx2 (1:100; Santa Cruz Biotechnology, sc-101145). Fins were washed in a high-salt 500 mM NaCl buffer for 30 min followed by extensive washes in PBS + 0.1% Tween-20. Secondary antibody incubations using Alexa Fluor conjugates (Thermo Fisher) were performed overnight protected from light at 4°C at a concentration of 1:1000 in blocking buffer. Fins were then washed extensively in PBS + 0.1% Tween-20, nuclei stained with Hoechst (Thermo Fisher) and mounted with SlowFade Diamond Antifade (Thermo Fisher).

### Paraffin section immunostaining

Dissected 32 dpf *shha:GFP* caudal fins were fixed in 4% PFA/PBS overnight at 4°C. After extensive PBS washing, fins were decalcified for 4 days in 0.5M EDTA, pH 8.0 with daily solution changes. Fins then were dehydrated in an ethanol series and tissue cleared with xylenes prior to longitudinal embedding in paraffin wax. 7 µm sections were cut on a Leica RM255 microtome. Antigen retrieval was performed on rehydrated sections using 1 mM EDTA + 0.1% Tween-20 for 5 minutes in a pressure cooker. Following PBS washes, sections were blocked in 1x PBS, 10% nonfat dry milk, 2% normal goat serum, and 4% fetal bovine serum for a minimum of 1 hour. Sections were incubated overnight at 4°C with primary antibodies in blocking solution. Primary antibodies were: anti-GFP (1:3000; AVES, GFP-1020), anti-Tp63 (1:100; Thermo Fisher, PA5-36039), anti-Laminin (1:40; Sigma, L9393), and anti-Zns5 (1:5, ZIRC). Sections were washed in PBS containing 500 mM NaCl + 0.1% Tween-20. Alexa Fluor conjugated secondary antibodies (Thermo Fisher) were diluted 1:1000 in blocking buffer and incubated for 1 hour at room temperature protected from light. Sections were washed, nuclei stained with Hoechst, and mounted with SlowFade Gold Antifade (Thermo Fisher). Images were acquired on a Zeiss LSM 880 laser scanning confocal microscope and images processed with Fiji-ImageJ, Imaris, and Adobe Photoshop.

### In vivo EdU incorporation assays

29 dpf *shha:GFP* juvenile fish were treated with DMSO or 1.25 µM BMS for 4 hours in groups of *n*=5. Anesthetized fish were injected intraperitoneally with 5 µl of 1 mg/mL EdU (Thermo Fisher) in sterile PBS, monitored for recovery for 10 minutes in fresh facility water, and then returned to treatment tanks. 12 hours post-injection, caudal fins were amputated and fixed for 4 hours at room temperature in 4% PFA/PBS. Fins were washed thoroughly with PBS and blocked overnight at 4°C in PBS/1% Triton X-100/5% Normal Donkey Serum/10% DMSO. EdU signal was detected with Click-iT Plus Alexa Fluor 647 Picolyl Azide (ThermoFisher) at 2.5 µl/ mL according to the manufacturer’s protocol. Following EdU detection, whole-mount GFP immunostaining and Hoechst nuclear staining was performed as described below. Whole mount confocal images were acquired using a Zeiss 880 LSM and 3D reconstructions prepared using Imaris. EdU+ and total intra-ray nuclei, i.e. from cells located in between the epidermal Shh domains of each hemi-ray, were identified and scored for Rays 2 and 3 using the Imaris “Spots” function and the following parameters: ROI around length of Shha:GFP+ domain, Quality Threshold 0.642, cell diameter 3 microns. Quantification of EdU+ cells is expressed as the number of EdU+ intra-ray cells over total number of Hoechst-stained nuclei.

### Cell migration imaging and analysis

Fish were anesthetized sequentially in freshly prepared 74 µg/ml tricaine solution for 3 minutes and monitored for slowed opercular movements. Anesthetized fish were transferred to a 35 mm FluroDish plate and mounted in 3% low melt agarose as described earlier. Set agarose was carefully removed from the most distal region of the caudal fin to allow for free movement of the epidermis while the trunk remained adhered to the FluoroDish. 74 µg/ml of tricaine solution was added to maintain anesthesia and cover the fin. After imaging, fish were returned to system water to confirm recovery and then promptly euthanized. Fish that did not recover were excluded from downstream cell migration analyses. We occasionally observed extremely rapid epidermal movements in which entire *shha:GFP+* domains would be shed from rays in <15 min. We suspect this phenomenon results from elevated stress, anesthesia intolerance, and/or damage from plate surface contact or agarose application. We excluded these animals from analyses.

For bulk cell migration assays, 22-24 dpf *ptch2:Kaede* fish were treated with 0.63 µM BMS or DMSO (*n*=8 per group). 24 hours later, fish were mounted and imaged with a Nikon Eclipse T*i*-E widefield microscope every 1 minute for 30 minutes. Imaris was used to automatically track cells for 3-6 single *ptch2:Kaede+* basal epidermal cells on dorsal rays 2-5 for each fish. All tracks were quality checked to confirm individual cell tracking. Individual cell speeds (arbitrary units, *n=*38 cells per group) and then average speed for each animal were determined. Statistical significance was assessed by Student’s unpaired *t*-tests for all cells tracked (*n=*38 cells per group) and average cell speed per fish (*n*=8 fish per group).

For position-dependent cell migration assays, 21-24 dpf *shha:GFP;runx2:mCherry* fish were treated with DMSO or 1.25 µM BMS for 24 hours prior to imaging. Fish were imaged every 2 minutes for 30 minutes with full z-stacks using a Nikon Ti2-E with a Yokogawa CSU-W1 SoRa spinning disk confocal unit. A single hemi-ray of Ray 2 or Ray 3 was analyzed for each time-lapse video. If both rays were captured, the ray with more pObs in frame was analyzed to avoid oversampling individuals. GFP+ cells were automatically tracked using Imaris software “Spots” algorithms with the following parameters: estimated cell diameter 5 microns, maximum distance between frames 6 microns, maximum gap between frames 3 time points. Each cell track was quality checked using 3D reconstructions and edited if Imaris assigned multiple cells to one track or fragmented the track of a given cell. 26-41 cells were tracked across 9 fish (*n=*5 for DMSO and *n*=4 for BMS groups, respectively) for a total of *n=*159 for DMSO-treated and *n*=135 for BMS-treated fish. Data was normalized for each fish by dividing the track speed of a single cell by the average of all cells tracked for that fish. Positional data was determined by setting the X-position of the most distal Runx2+ pOb as “0” and assigning a relative initial X-position for each *shha:GFP+* cell. Cells with a negative starting position were therefore pOb-associated while those with a positive starting position had already migrated beyond the pOb pool when video acquisition began. Imaris was used to determine each cell’s total X-displacement and track speed. Graphs were generated using GraphPad Prism V8. Fourth-order best-fit polynomial curves were added to position/speed graphs to help visualize data trends.

## ACKNOWLEDGEMENTS

We thank the University of Oregon AqACS Facility for zebrafish care; the University of Oregon zebrafish community for support; G. Crump for providing the *runx2:mCherry* fish generated by S. Fisher’s group; and C. Kimmel, A. Saera-Vila, and the Stankunas lab for input.

## FOOTNOTES

### Author contributions

J.A.B, A.E.R. and K.S. designed experiments with input from S.S.; J.A.B and A.E.R. performed experiments; J.A.B, A.E.R. and K.S. prepared and wrote the manuscript.

## Funding

The National Institutes of Health (NIH) provided research funding (1R01GM127761 (K. S. and S. S.). J.A.B. had an Experiencing Science Practices through Research to Inspire Teaching (ESPRIT) award supported by the National Science Foundation’s Robert Noyce Teacher Scholarship Program, a Mary G. Alden Scholarship, and an Institute of Molecular Biology Summer Scholarship. A.E.R. received support from the University of Oregon Genetics Training Program (5T32GM007413) and a NIH NRSA fellowship (1F31GM139343).

### Data and material availability

Requests for materials should be addressed to K. S.

## SUPPLEMENTARY MATERIAL

**Figure S1.**
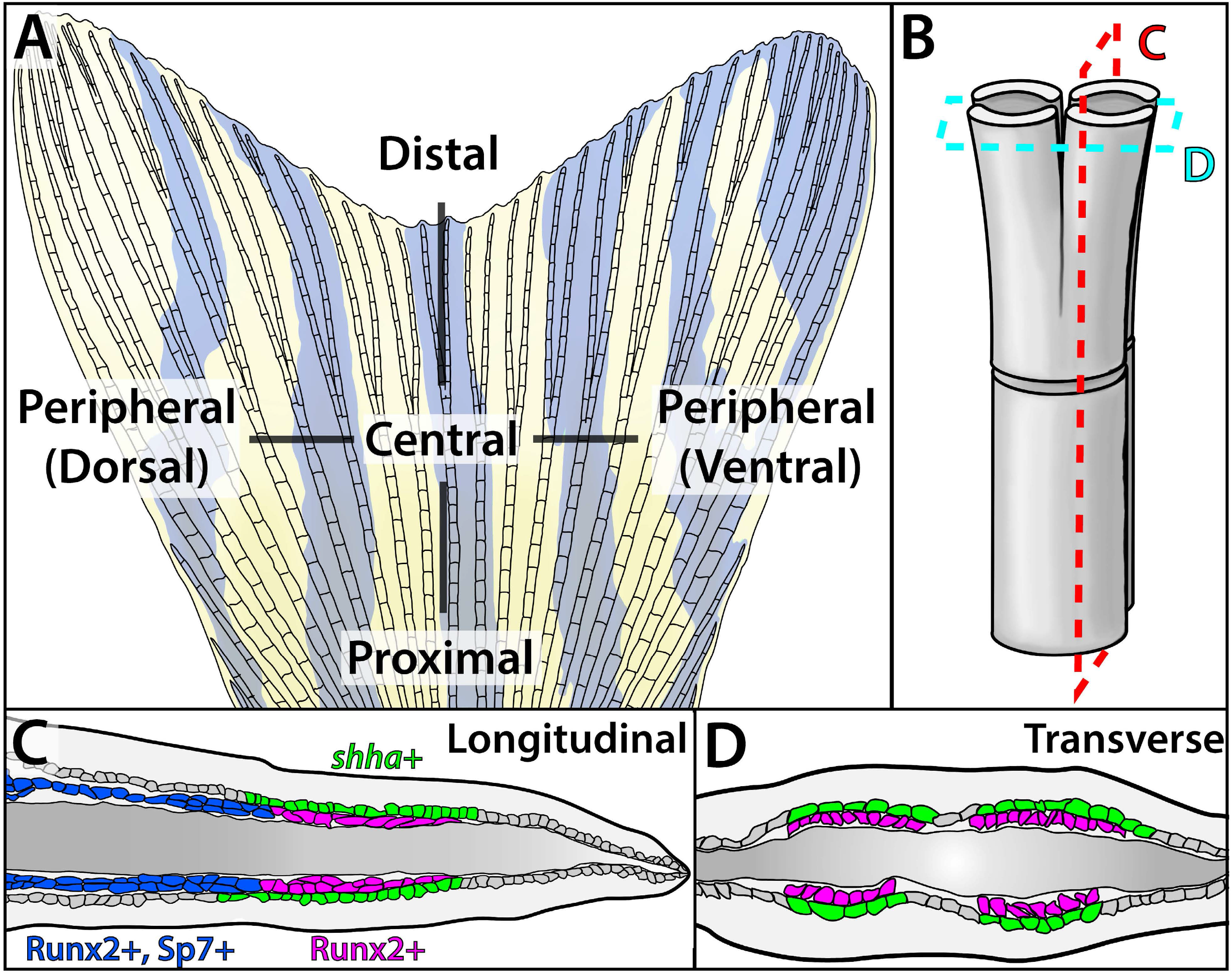
Schematic of caudal fin skeletal anatomy. (A) Schematic of an adult zebrafish caudal fin. The caudal fin skeleton comprises 18 bony rays (lepidotrichia), of which the inner 16 branch at least once. (B) Cartoon rendering of a skeletal ray branch point. Two opposed hemi-cylindrical calcified hemi-rays form each ray. Branching produces two equally sized daughter rays. (C, D) Colored tracings of longitudinal (C) and transverse (D) sections through a branching lepidotrichia. Distal domains of *shha*-expressing basal epidermal cells (basal epidermis, green) directly neighbor Runx2+ pre-osteoblasts (magenta). Those basal epidermal cells distally beyond pre-osteoblast pools lose *shha* expression. *shha*+ basal epidermis and pre-osteoblast domains both split peripherally as branching initiates. As the fin extends, the distal Runx2+ pre-osteoblast pool generates differentiating Runx2/Sp7+ osteoblasts (purple) that eventually mature into proximal sp7+ bone-forming cells (blue).

**Figure S2.**
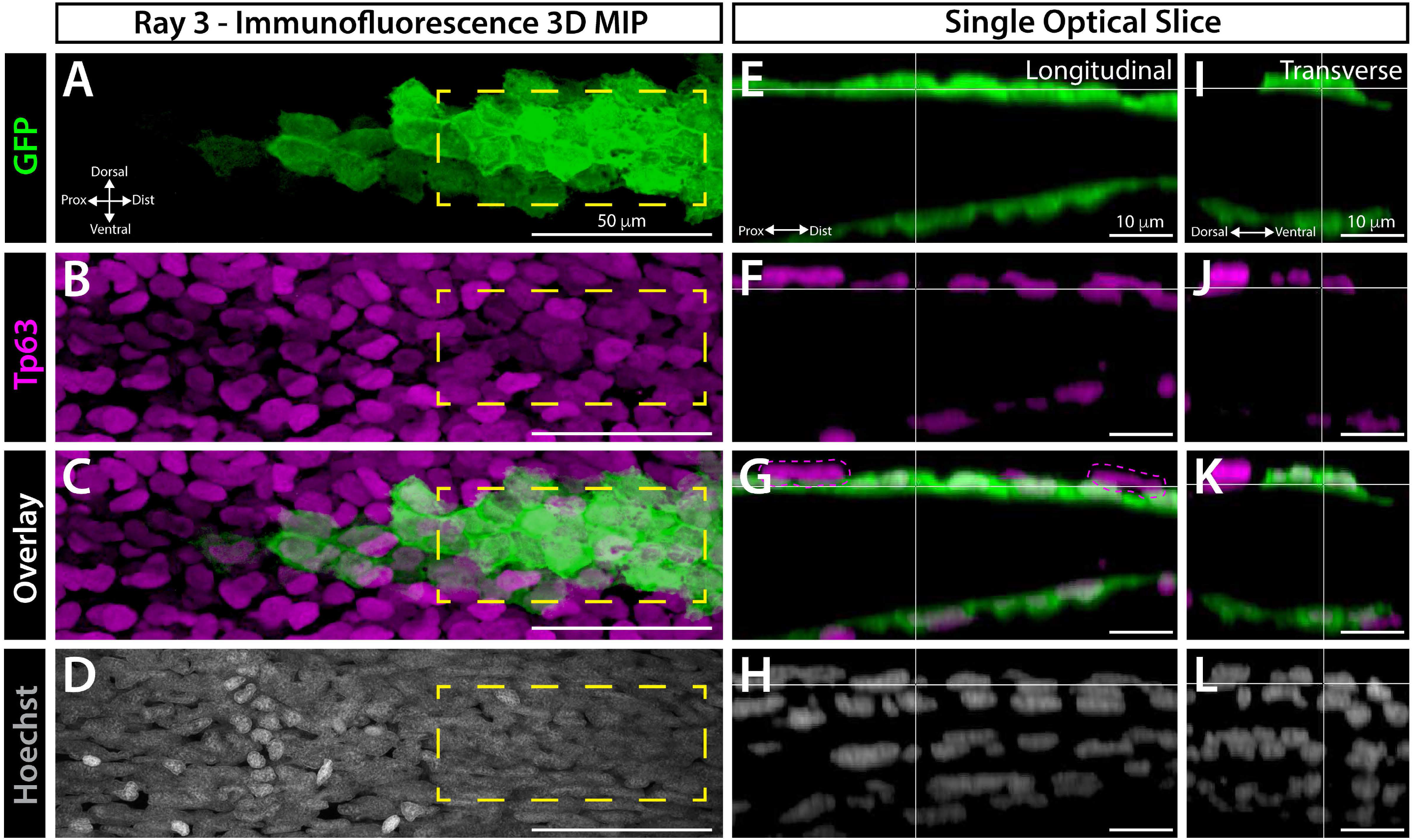
*shha* is expressed in a single layer of distal basal epidermal cells in developing caudal fins. 3D reconstructed sectional views of dorsal ray 3 from a whole mount immunostained 23 dpf *shha:GFP* caudal fin. Panels show GFP (green, A, E, I), the basal epidermal marker Tp63 (magenta, B, F, J), GFP and Tp63 overlays (C, G, K; G and K are reproduced in Figure 1), and Hoechst-stained nuclei (white, D, H, I). (A-D) Maximum intensity projection (MIP) of the “native” frontal view. (E-L) Reconstructed single optical slice equivalents showing longitudinal (E-H) and transverse (I-L) planar views of the dashed yellow boxed regions in (A-D). Shha-expressing cells (as marked by GFP) define the innermost basal epidermal cell layer and all co-express Tp63. An occasional single-positive Tp63+ basal epidermis (representative cell in magenta dashed lines, G) is found in a second, outer layer of basal epidermis. Scale bars and orientations are indicated.

**Figure S3.**
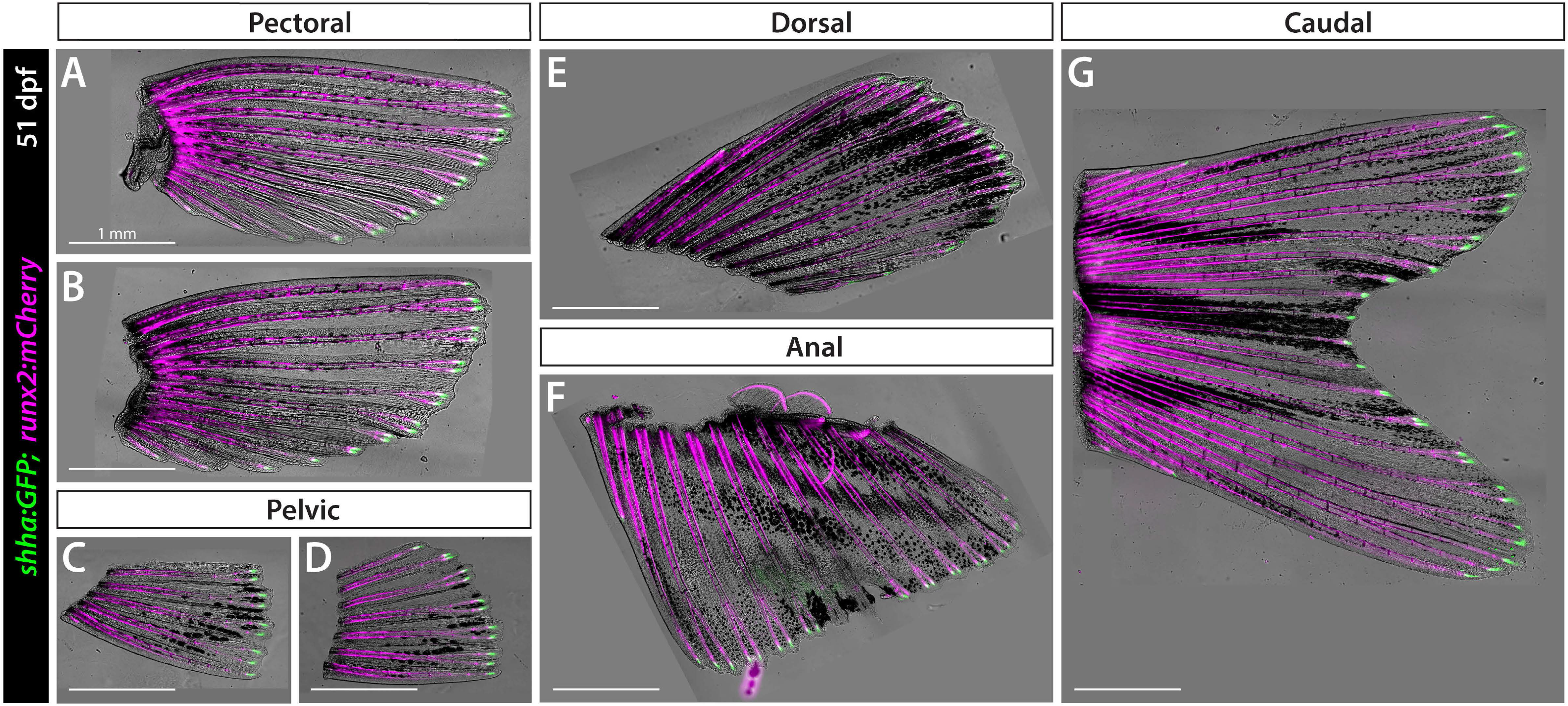
Distal *shha* expression is conserved across developing fins. Dissected fins from a representative 51 dpf *shha:GFP;runx2:mCherry* fish. Zebrafish have 7 fin appendages: the paired pectoral (A, B) and pelvic (C, D) fins and unpaired dorsal (E), anal (F), and caudal (F) fins. All have branched rays marked by *runx2:mCherry* (magenta) with *shha:GFP+* domains (green) at the distal end of each developing ray. All images are brightfield/GFP/mCherry overlays and sized to the same scale (1 mm scale bars).

**Figure S4.**
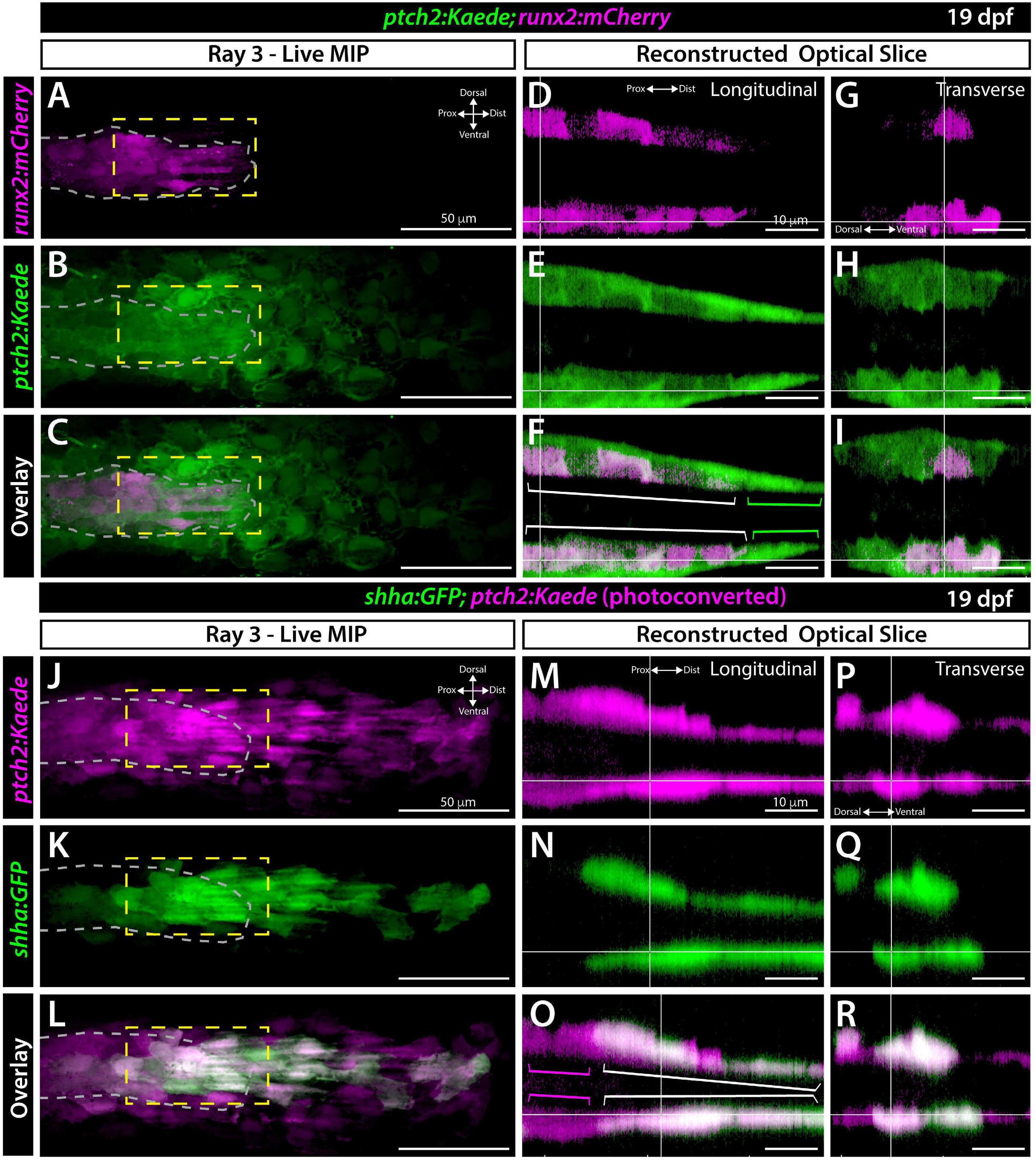
*ptch2* is expressed in tightly associated layers of distal basal epidermal cells and pre-osteoblasts in developing caudal fins. (A-I) Whole mount confocal imaging of dorsal ray 3 from a live 19 dpf *ptch2:Kaede; runx2:mCherry* caudal fin. (A-C) Maximum intensity projection (MIP) of a frontal view. The pre-osteoblast pool is outlined with grey dashed lines. *runx2:mCherry*-marked pre-osteoblasts are in magenta (A) and *ptch2:Kaede* is in green (B). The overlay is shown in (C). (D-I) Reconstructed optical slice views of the region marked by yellow dashed lines in A-C. Overlays (F, I; reproduced in main Figure 1) show relatively proximal regions of *ptch2:Kaede* and *runx2:mCherry+* co-expressing pre-osteoblasts (white brackets) and an adjacent thin layer of *ptch2:Kaede+* basal epidermal cells. These basal epidermal cells extend further distally from the pre-osteoblast pool (green brackets). (J-R) Confocal images of dorsal ray 3 from a 19 dpf *shha:GFP;ptch2:Kaede* caudal fin in which the Kaede protein has been photoconverted from green to red fluorescence emission. (J-L) Frontal view MIP with osteoblast-populated region outlined with grey dashed lines. *shha:GFP* basal epidermis (green) co-express *ptch2:Kaede* (magenta). (M-R) Reconstructed optical slices of the yellow dashed boxes in J-L. Overlays (O, R; reproduced in main Figure 1) demonstrate proximal regions contain single-positive *ptch2:Kaede+* pre-osteoblasts (magenta brackets) whereas distal regions include co-expressing *shha:GFP* and *ptch2:Kaede* basal epidermis (white brackets). Scale bars are 50 μm in A-C, J-L and 10 μm in D-I, M-R.

**Figure S5.**
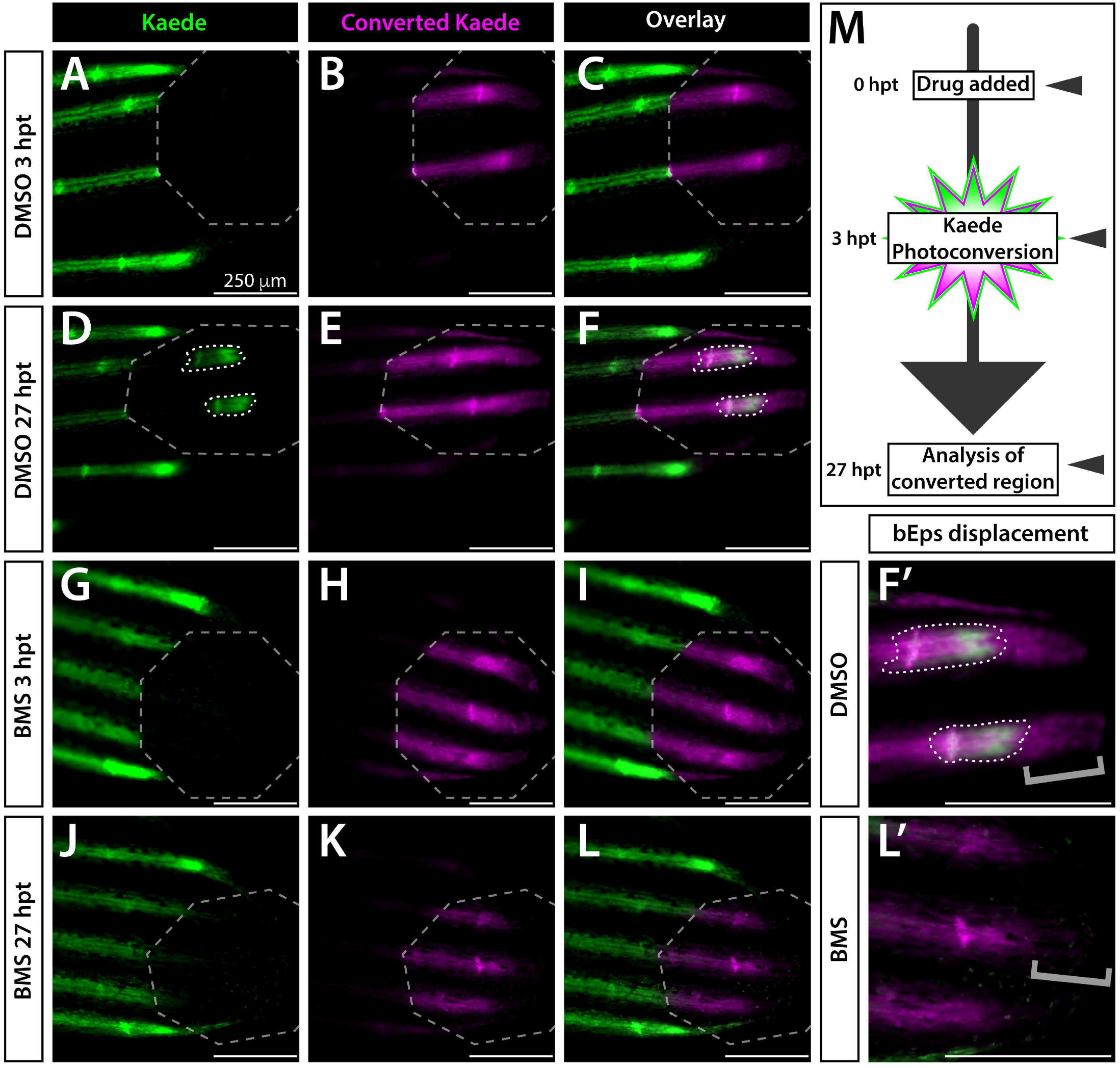
BMS-833923 inhibits Shh/Smo signaling in developing caudal fins. Expanded data from Figure 2. (A-L) Whole mount fluorescence images of the distal caudal fin of 25 dpf *ptch2:Kaede* fish treated with DMSO (A-F) or BMS-833923 (BMS; G-L), imaged at the time of Kaede photoconversion (3 hours post treatment (hpt)) and 24 hours later (27 hpt). Grey dashed octagons mark the photoconverted regions of interest (ROIs). BMS exposure prevented production of new green fluorescent Kaede within the ROI (J-L). (F’, L’) Zoom view of distal ray regions. Brackets mark presence or absence of photoconverted basal epidermal cells that migrated distally over the 24 hour “chase”. These cells are missing in BMS-treated fish (L’), likely due to accelerated movement and therefore shedding. The few green Kaede+ basal epidermal cells in (J, L) moved into the photoconverted region without producing new Kaede. (M) Schematic of the time course for drug treatments, photoconversion, and imaging. Imaged fish represent groups of *n*=8. Scale bars are 250 μm.

**Figure S6.**
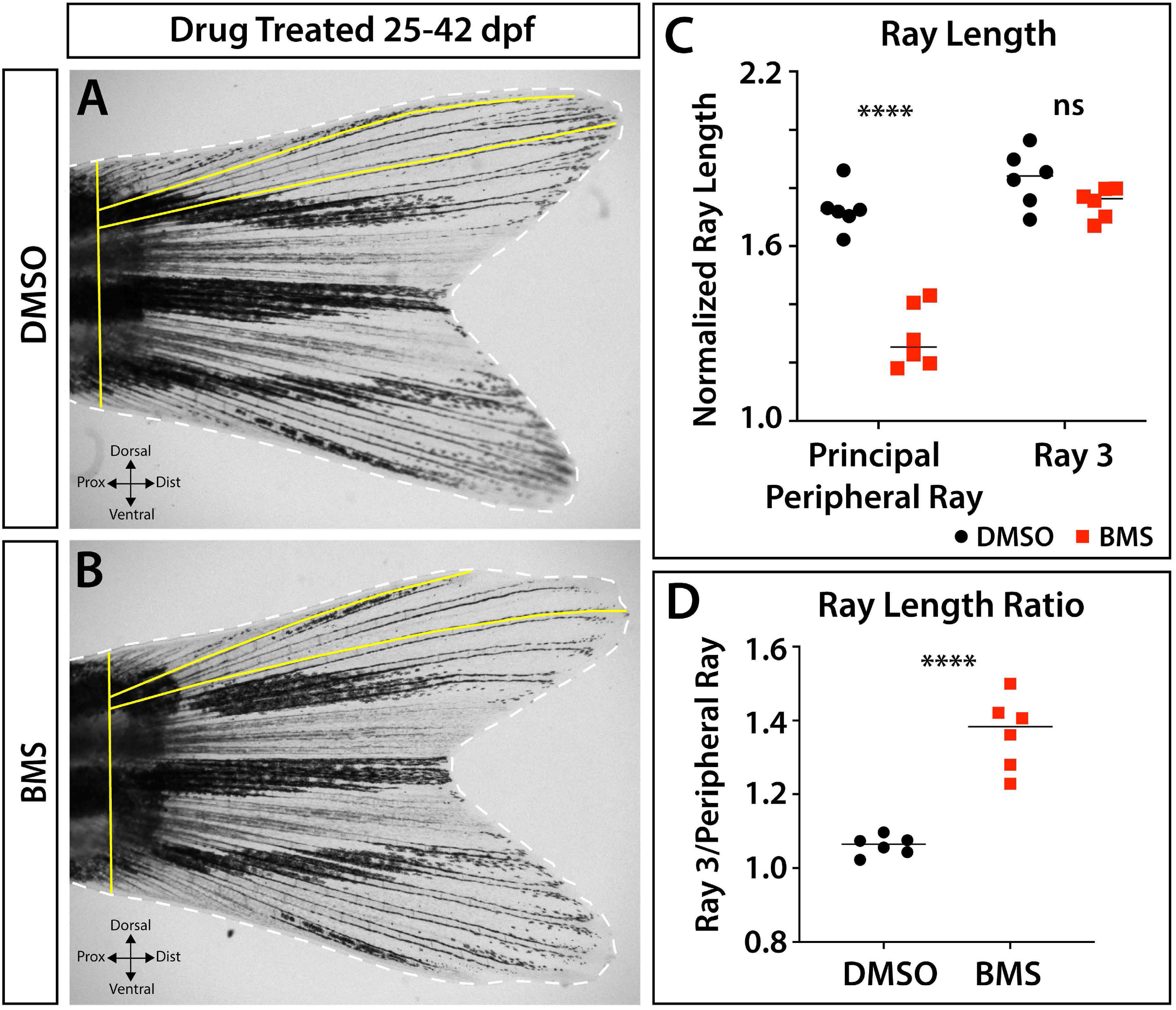
Shh/Smo signaling contributes to principal peripheral ray outgrowth. Whole mount caudal fin images of (A) DMSO- and (B) BMS-treated juvenile fish (exposed from 25-42 days post fertilization (dpf)) and (C) graphs of ray morphometrics. Yellow lines indicate measured ray lengths and fin widths. Dorsal ray 3 does not significantly differ in length between treatment groups (*p* = 0.09). In contrast, the non-branching principal peripheral ray 1 is uniquely shorter in BMS-treated versus DMSO control fish (*p* < 0.0001). As such, the ray length ratio between ray 3 and principal peripheral ray 1 is higher in BMS-treated fish (*p* < 0.0001). Images are representatives of experimental groups of *n=*6. Ray length measurements are from the base of the fin and are normalized to fin width, which did not differ between groups. Statistical significance was determined using Student’s unpaired t-tests.

**Figure S7.**
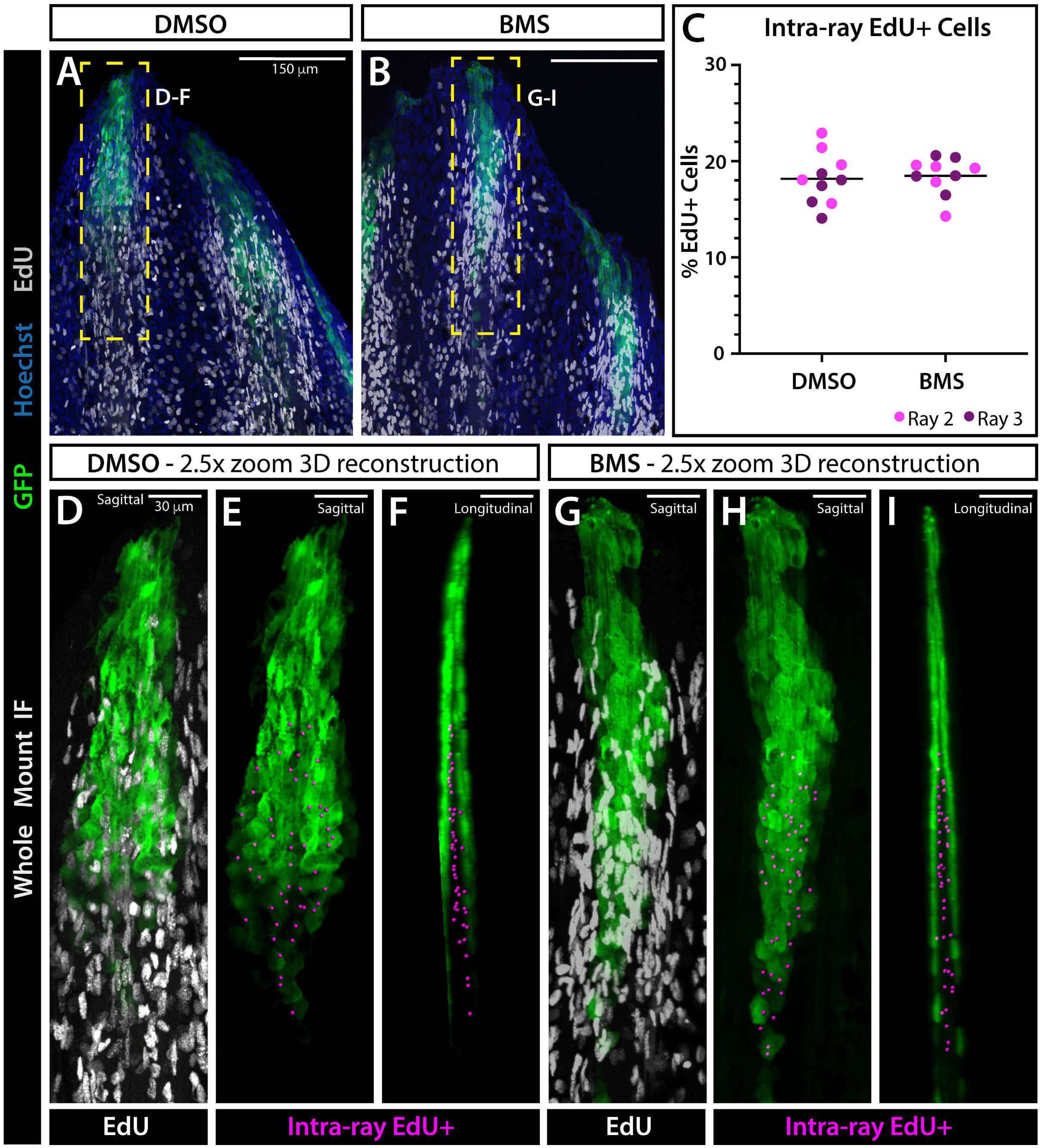
Shh/Smo signaling does not impact proliferation in developing fin rays. (A, B) Confocal maximum intensity projection images of whole mount immunostained 29 dpf developing *shha:GFP* caudal fins. Fish were treated with DMSO (A) or BMS-833923 (BMS) (B) for 4 hours, IP injected with EdU, returned to drug, and fins harvested 12 hours later. GFP is in green, Hoechst nuclear stain in blue, and EdU as detected with Click-iT Plus in white. (C) Scatter plot graph showing the percent of intra-ray EdU+ cells, i.e. those located in between the epidermal Shh domains of each hemi-ray, does not significantly differ between DMSO controls and BMS-treated fish. *n=*5 for each group. (D-I) 3D reconstructions of ray 3 domains marked by yellow dashed boxes in (A, B). (D, G) Overlay of GFP and EdU, sagittal view. (E, H) GFP with magenta spheres marking EdU+ cells located within the intra-ray space, detected and quantified with Imaris software. (F, I) Longitudinal view of (E, H). Intra-ray EdU+ cells are located in between Shh+ domains with epidermal cells excluded from automated scoring. Scale bars are 150 μm or 30 μm, as indicated.

**Figure S8.**
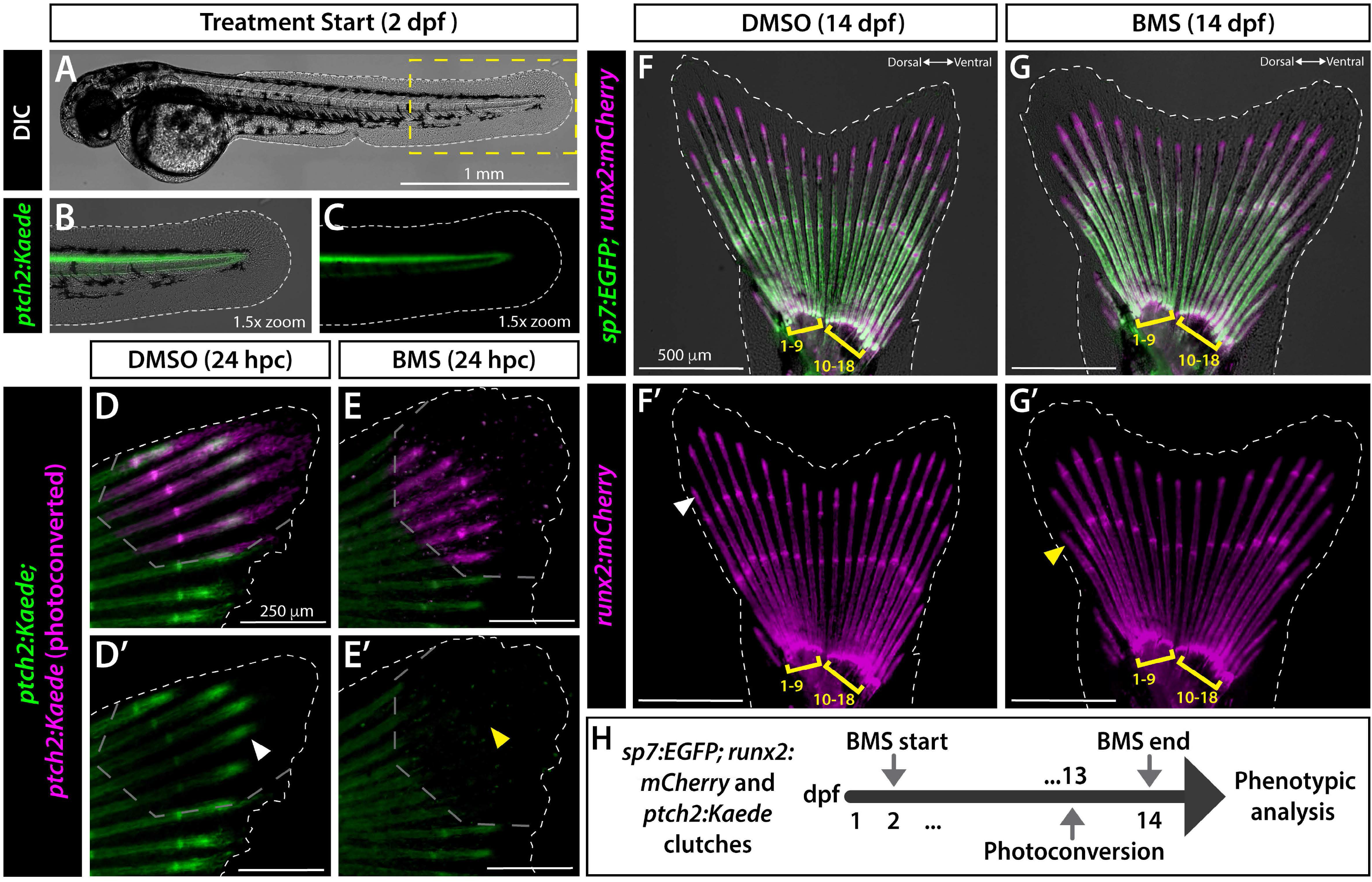
Shh/Smo signaling does not influence initial caudal fin patterning. (A-C) Whole mount differential interference contrast (DIC) and/or fluorescence images of a 2 dpf *ptch2:Kaede* embryo. The dashed yellow box marks the 1.5x zoomed region shown in (B, C). The primordial fin fold lacks rays or structural ray precursors. *ptch2:Kaede* expression is restricted to the notochord and does not expand into the fin fold. (D-E’). *ptch2:Kaede* larvae treated with DMSO or BMS-833923 (BMS) from 2 dpf, photoconverted distal fin regions at 13 dpf (grey dashed lines), and imaged 24 hours later (*n*=3-5 per group). BMS treatment blocked production of new green Kaede (yellow arrowhead, E’) compared to controls (white arrowhead, D’). (F-G’) Whole mount fluorescence images of *sp7:EGFP;runx2:mCherry* larvae treated with DMSO (F) or BMS (G) from 2-14 dpf. Both groups have caudal fins with the standard complement of 18 segmented rays, with 9 rays each on the dorsal and ventral lobes (yellow brackets). BMS-treated fins exhibit shortened principal peripheral rays (yellow arrowhead, G’) compared to DMSO controls (white arrowhead, F’). *n*=33 DMSO- and *n*=44 BMS-treated larvae from 2-14 dpf. Scale bars are 1 mm, 250 μm, or 500 μm, as marked.

**Figure S9.**
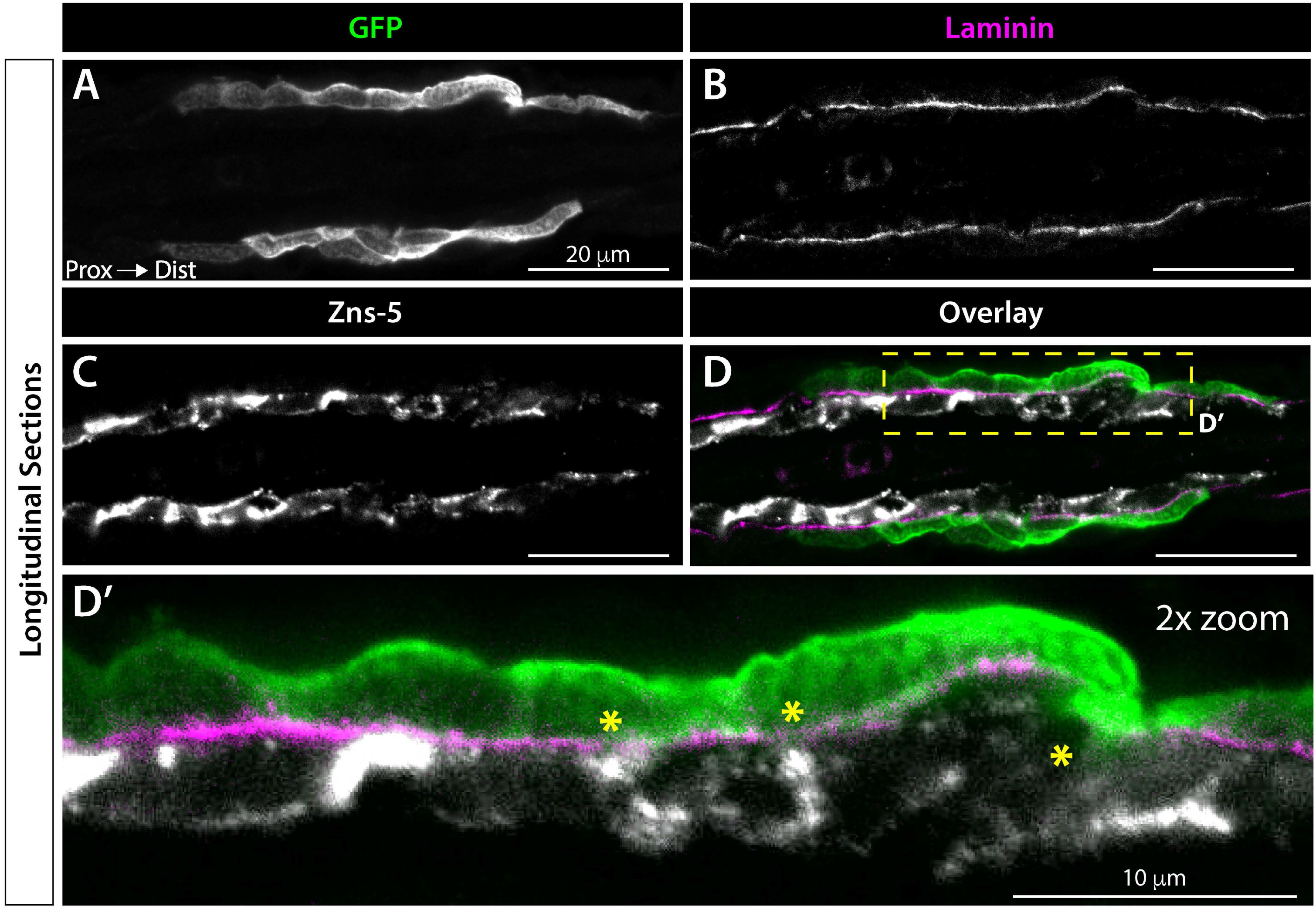
Shha+ basal epidermis are tightly associated with pre-osteoblasts in distal regions of incomplete basement membrane assembly. (A-D) Confocal images of the distal ray regions from immunostained longitudinal caudal fin sections of 32 dpf *shha:GFP* fish. GFP-expressing basal epidermal cells (green) and Zns-5-marked osteoblasts (white) are separated by a Laminin-defined basement membrane (BM; magenta). (D’) 2x magnification of the yellow dashed box region in the (D) overlay. Asterisks mark areas where Laminin signal is less dense, indicating an incompletely assembled BM, and GFP and Zns-5 partially overlap (D’). Scale bars are 20 μm.

**Figure S10.**
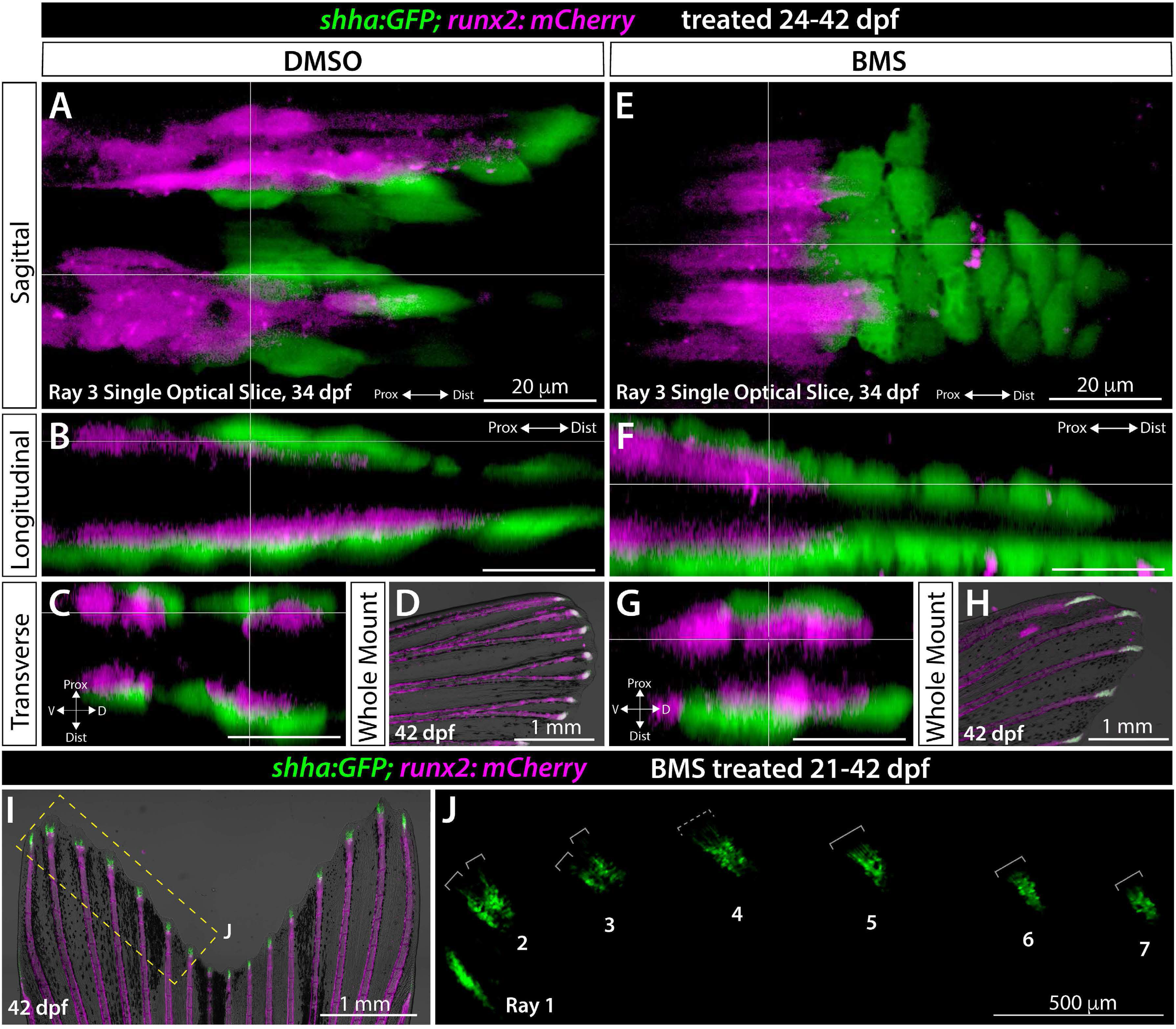
Shh/Smo signaling does not influence Shh+ basal epidermis and pre-osteoblast juxtaposition. (A-J) Confocal or widefield fluorescence caudal fin images from *shha:GFP;runx2:mCherry* fish treated with DMSO (control) or 1.25 μM BMS-833923 (BMS) starting at 24 dpf (*n*=8 per group). (A-C, E-G) Single optical slices or equivalent 3D-reconstructed views of 34 dpf mid-treatment fish, imaged concurrently with active ray branching morphogenesis. *shha:GFP+* basal epidermal cells (green, A) are closely associated with *runx2:mCherry*-expressing pre-osteoblasts (magenta). BMS-treated fish (E-G) maintain close Shh+ basal epidermal and Runx2+ pre-osteoblast contacts while lacking clear *shha:GFP* domain-splitting (*n*=4 per group). (D, H, I, J) Caudal fin images of *shha:GFP;runx2:mCherry* fish continued on drug treatment until 42 dpf (*n*=4 per group). (D, H, I) Widefield fluorescence and brightfield overlay caudal fin images. Unlike DMSO controls (D), BMS-treated fish do not develop branched rays, confirming drug efficacy (H). (I, J) Fin of a 21-42 dpf BMS-treated fish. The yellow dashed box indicates the GFP-alone magnified panel in (J). *shha:GFP* domains are variably split (two grey br ets, rays 2, 3), partially split (dashed grey bracket, ray 4) or unsplit (one grey bracket, rays 5-7). Scale bars are 20 μm (A-C, E-G), 1 mm (G, H, I) and 500 μm (J).

**Figure S11.**
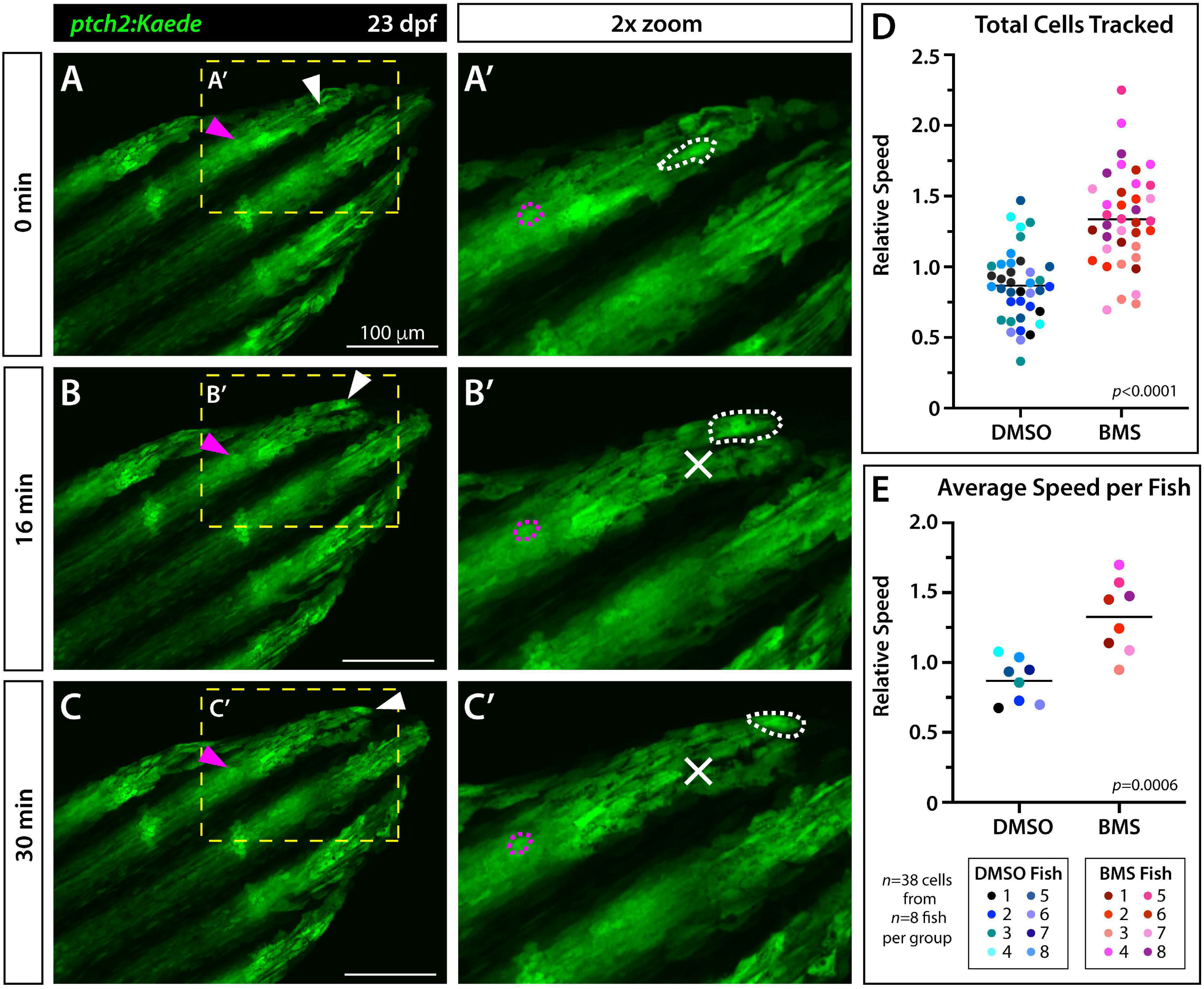
Shh/Smo signaling restrains the distal migration of basal epidermal cells. (A-C) Still frames from 30 minute time lapse movies capturing basal epidermal distal migration in a caudal fin of a live mounted 23 dpf *ptch2:Kaede* fish. The fish was DMSO-treated as a control group member for quantitative studies. Maximum intensity projections of a fin’s dorsal lobe are shown at 0 minutes (A, start position), 16 minutes (B, halfway through video), and 30 minutes (C, end position). White and magenta arrowheads indicate a *ptch2:Kaede*+ basal epidermis and pre-osteoblast, respectively. Yellow dashed boxes mark the 3x magnified distal ray region in (A’-C’). The representative basal epidermis (white) and pre-osteoblast (magenta) are outlined with dashed lines. The white Xs in (B’, C’) indicate the basal epidermal cell’s starting position at 0 minutes. Scale bars are 100 μm. (D) Scatterplot graph showing average speeds (arbitrary units) of individual *ptch2:Kaede*+ basal epidermal cells from fish treated with DMSO or BMS-833923 (BMS) for 24 hours and then imaged over 30 minutes. Cells of BMS-treated fish migrate faster than DMSO controls (*p* < 0.0001, Student’s t-test). (E) Graph showing basal epidermal migration averaged by individual animal (as indicated by different colors) is significantly faster (*p* < 0.0006, Student’s t-test) in BMS-treated fish. *n*=3-6 cells per fish with 8 fish per treatment for a total of 38 tracked cells per group. Colors represent single basal epidermal cells from the same animal.

**Figure S12.**
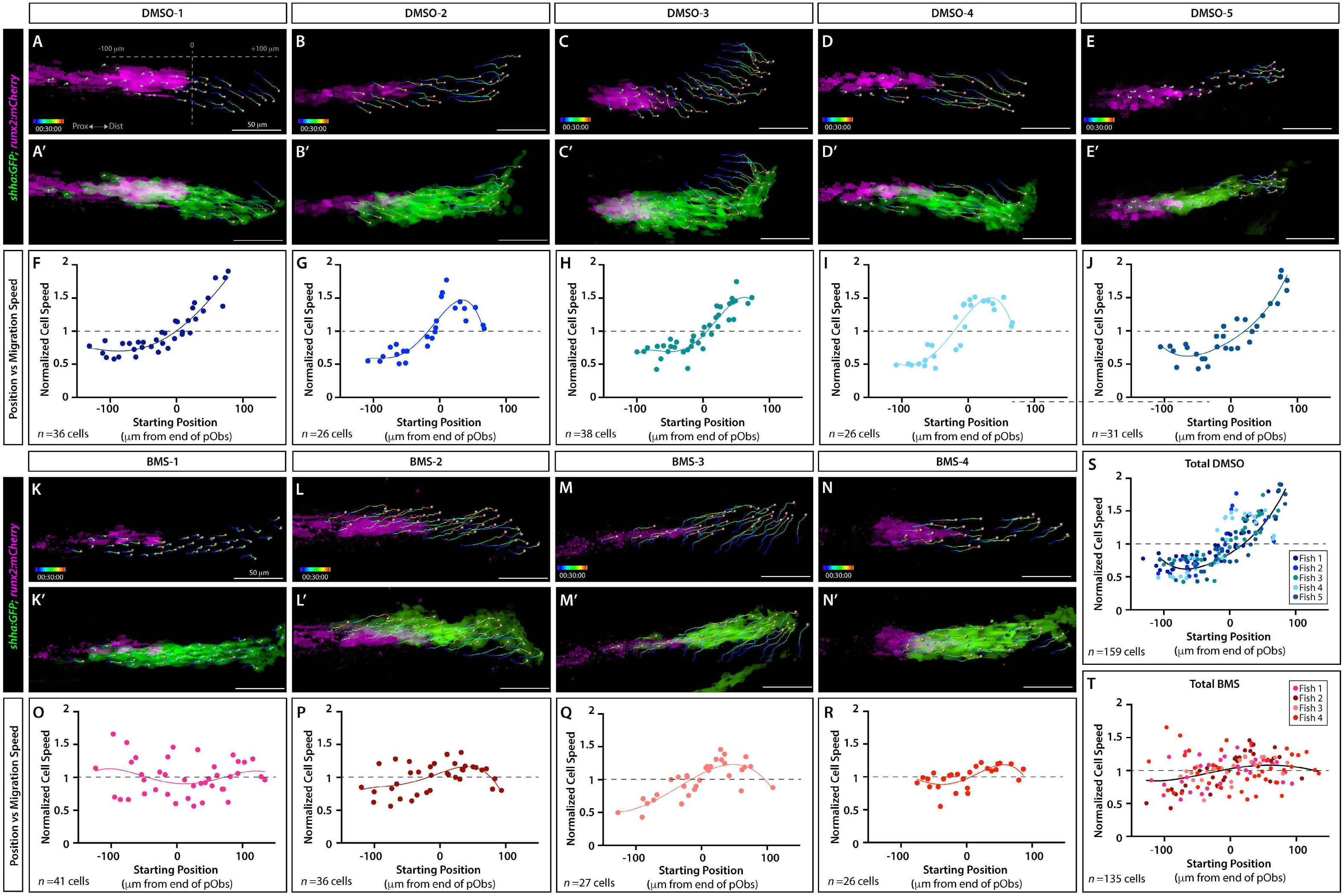
Position- and Shh/Smo-dependent *shha:GFP+* basal epidermal migration rates of individual fish. Expanded data from Figure 6. (A-E’, K-N’) Maximum intensity projections of the final 30 minute frame of time lapse-imaged *shha:GFP;runx2:mCherry* fish (21-24 dpf) treated with DMSO (*n*=5) or BMS-833923 (BMS; *n*=4) for 24 hours. Panels show *runx2:mCherry* pre-osteoblasts only (A-E, K-N; magenta) or both pre-osteoblasts and *shha:GFP+* basal epidermal cells (A’-E’, K’-N’; green). Grey spheres mark semi-automatically tracked basal epidermal cells with their cell displacement over 30 minutes indicated by multicolor tracks. (F-J, O-T) Average speed vs. starting position scatterplot graphs for each fish. Data points represent individual cell speeds. Non-parametric best-fit curves provide a visual guide. (S, T) Summary graphs reproduced from Figure 6 show combined data with added overall trends. Scale bars are 50 μm.

**Figure S13.**
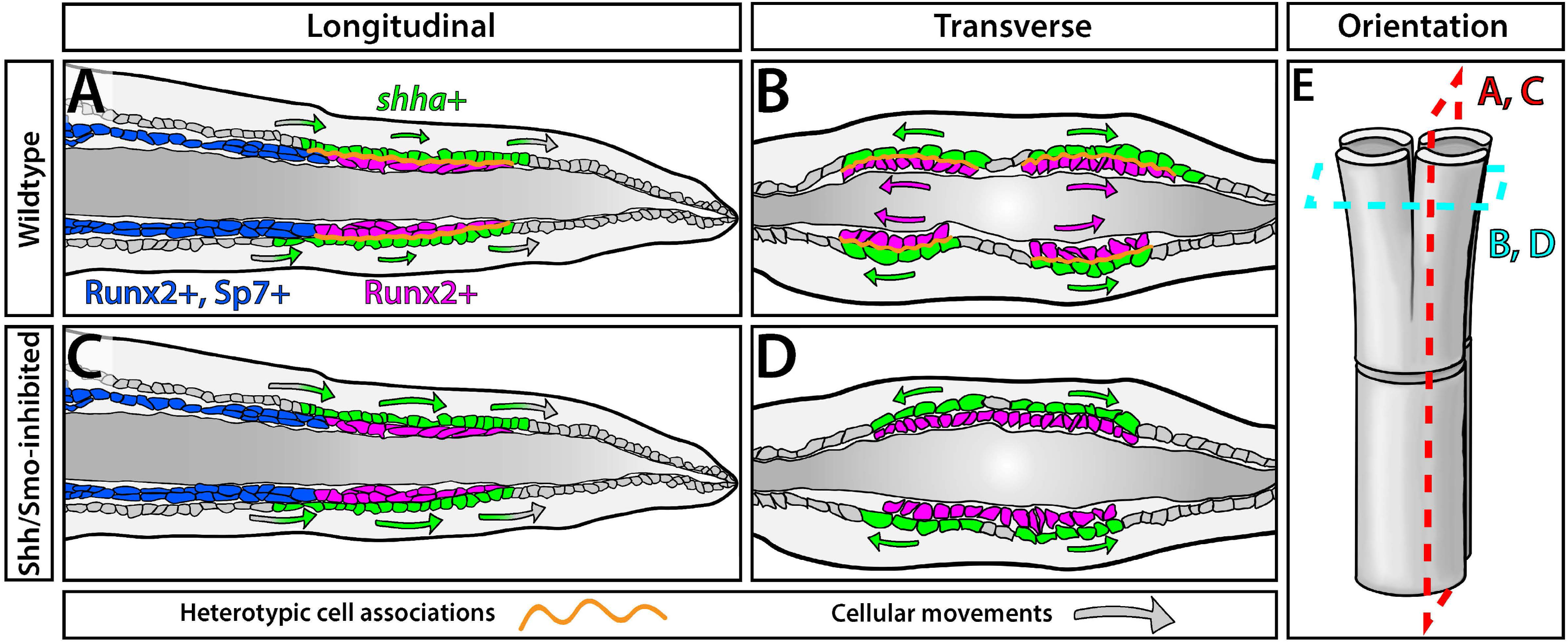
Basal epidermis collective movements and Shh/Smo-driven heterotypic associations with pre-osteoblasts direct fin ray branching morphogenesis. (A-D) Schematics depicting the organization and cell movements of basal epidermal cells and osteoblasts during ray branching of outgrowing fins. Wildtype (A, B) and Shh/Smo-inhibited (C, D) conditions are shown. Longitudinal views (A, C) illustrate basal epidermal distal cell movements and transverse views (B, D) highlight lateral movements of both basal epidermal cells and Runx2+ pre- osteoblasts. (E) Schematic of a mature branched ray to orient the equivalent sectional views in (A-D). (A, C) Distal-moving basal epidermal cells (grey) enter the Shh/Smo active zone and upregulate *shha* to activate a Shh/Smo response in themselves and juxtaposed pre-osteoblasts. Shh/Smo-inhibition using the small molecule BMS-833923 increases and steadies the rate of basal epidermal distal migration (see animation in Movie 5). This suggests Shh/Smo activity normally slows the distal march of transiently Shh-expressing basal epidermal cells by enhancing cell associations (orange lines) with relatively static Runx2+ pre-osteoblasts. Basal epidermal cells downregulate *shha* and lose their Shh/Smo response as they move beyond the pre-osteoblasts. (B, D) The *shha-*expressing basal epidermal domain laterally divides, closely followed by underlying pre-osteoblast pools, prior to ray branching. BMS-833923 prevents pre-osteoblasts from following the lateral-splitting *shha*-expressing basal epidermis and thereby blocks ray branching. Shh/Smo-promoted heterotypic cell associations may transfer lateral forces from successively passaging basal epidermal cells to progressively re-position pre-osteoblasts into split pools. Each pre-osteoblast pool and their derived Runx2+/sp7+ osteoblasts now form separate daughter rays joined at a branch point.

**Movie 1.** A dynamic 3-D visualization of confocal-imaged caudal fin dorsal ray 3 from a 28 dpf *shha:GFP;runx2:mCherry* fish. *shha:GFP+* basal epidermis (green) of both hemi-rays are beginning to branch. Underlying pre-osteoblasts (magenta) are pressed against basal epidermis. 3D reconstruction of live confocal microscopy. Surfaces added with Imaris to visualize domain boundaries.

**Movie 2.** 3-D space exploration of the distal region of a confocal-imaged caudal fin hemi-ray from a 28 dpf *shha:GFP;runx2:mCherry* fish. The *shha:GFP+* basal epidermis (green) domain has started splitting ahead of ray branching. A ridge of pre-osteoblasts (magenta) is nestled in a hollow of *shha:GFP+* basal epidermis. Imaris-generated surfaces help visualize domain boundaries. Single optical slices of the same region are shown in Figure 5.

**Movie 3.** Time lapse movie of the dorsal lobe of the caudal fin of a live-mounted 23 dpf *ptch2:Kaede* fish treated with DMSO (control group) for 24 hours prior to imaging. Widefield fluorescence images were collected every 2 minutes over a 30-minute period. Still images and data analysis are in Figure S11.

**Movie 4.** Time lapse movie of caudal fin dorsal lobes of representative live-mounted *shha:GFP;runx2:mCherry* fish treated with DMSO (upper panel) or BMS-833923 (lower panel) for 24 hours prior to imaging. Shh+ basal epidermis in green and Runx2+ pre-osteoblasts in magenta were imaged by fluorescence confocal microscopy. Multicolor lines indicate cell migration tracks for individual Shh+ basal epidermal cells (marked by grey spheres) over time. Images were acquired every 2 minutes for 30 minutes. Still images and data analysis are in Figure 6.

**Movie 5.** Heterotypic cell associations decrease the rate of basal epidermal distal movements while adjacent to pre-osteoblasts. This animation related to Figure S13 highlights the distal transit of individual basal epidermal cells (red) in longitudinal section views of outgrowing caudal fins. (A) Basal epidermal cells constantly and collectively move distally as an interconnected epithelium, upregulating *shha* to sustain a distal Shh/Smo active zone including directly adjacent pre-osteoblasts. Normally, Shh/Smo-promoted associations between basal epidermal cells and pre-osteoblasts transiently restrain the passage of basal epidermal cells. Basal epidermal cells then accelerate once passing beyond the pre-osteoblasts. (B) Shh/Smo inhibition using BMS-833923 tempers associations between bEps and pObs to increase and make constant the rate of basal epidermis distal movement. The Shh/Smo-dependent heterotypic interactions may transfer basal epidermal lateral movement forces to progressively split pre-osteoblast pools for ray branching morphogenesis.

## REFERENCES

Ahn, Y., Sanderson, B.W., Klein, O.D., Krumlauf, .R, 2010. Inhibition of Wnt signaling by Wise (Sostdc1) and negative feedback from Shh controls tooth number and patterning. Development 137, 3221–3231. https://doi.org/10.1242/dev.054668

Akimenko, M.-A., Ekker, M., 1995. Anterior Duplication of the Sonic hedgehog Expression Pattern in the Pectoral Fin Buds of Zebrafish Treated with Retinoic Acid. Developmental Biology 170, 243–247. https://doi.org/10.1006/dbio.1995.1211

Alexandre, C., Jacinto, A., Ingham, P.W., 1996. Transcriptional activation of hedgehog target genes in Drosophila is mediated directly by the cubitus interruptus protein, a member of the GLI family of zinc finger DNA-binding proteins. Genes & Development 10, 2003–2013. https://doi.org/10.1101/gad.10.16.2003

Aman, A.J., Fulbright, A.N., Parichy, D.M., 2018. Wnt/β-catenin regulates an ancient signaling network during zebrafish scale development. eLife 7, e37001. https://doi.org/10.7554/elife.37001

Anderson, E., Peluso, S., Lettice, L.A., Hill, R.E., 2012. Human limb abnormalities caused by disruption of hedgehog signaling. Trends in Genetics 28, 364–373. https://doi.org/10.1016/j.tig.2012.03.012

Armstrong, B.E., Henner, A., Stewart, S., Stankunas, K., 2017. Shh promotes direct interactions between epidermal cells and osteoblast progenitors to shape regenerated zebrafish bone. Development 144, 1165–1176. https://doi.org/10.1242/dev.143792

Ayers, K.L., Gallet, A., Staccini-Lavenant, L., Thérond, P.P., 2010. The Long-Range Activity of Hedgehog Is Regulated in the Apical Extracellular Space by the Glypican Dally and the Hydrolase Notum. Developmental Cell 18, 605–620. https://doi.org/10.1016/j.devcel.2010.02.015

Barske, L., Fabian, P., Hirschberger, C., Jandzik, D., Square, T., Xu, P., Nelson, N., Yu, H.V., Medeiros, D.M., Gillis, J.A., Crump, J.G., 2020. Evolution of vertebrate gill covers via shifts in an ancient Pou3f3 enhancer. Proceedings of the National Academy of Sciences 117, 24876–24884. https://doi.org/10.1073/pnas.2011531117 PMID - 32958671

Bellusci, S., Furuta, Y., Rush, M.G., Henderson, R., Winnier, G., Hogan, B.L., 1997. Involvement of Sonic hedgehog (Shh) in mouse embryonic lung growth and morphogenesis. Development (Cambridge, England) 124, 53–63.

Chen, C.-H., Puliafito, A., Cox, B.D., Primo, L., Fang, Y., di Talia, S., Poss, K.D., 2016. Multicolor Cell Barcoding Technology for Long-Term Surveillance of Epithelial Regeneration in Zebrafish. Developmental Cell 36, 668–680. https://doi.org/10.1016/j.devcel.2016.02.017

Chen, J.-S., Huang, X., Wang, Q., Huang, J.-Q., Zhang, L., Chen, X.-L., Lei, J., Cheng, Z.-X., 2013. Sonic hedgehog signaling pathway induces cell migration and invasion through focal adhesion kinase/AKT signaling-mediated activation of matrix metalloproteinase (MMP)-2 and MMP-9 in liver cancer. Carcinogenesis 34, 10–19. https://doi.org/10.1093/carcin/bgs274

Chen, Q., Xu, R., Zeng, C., Lu, Q., Huang, D., Shi, C., Zhang, W., Deng, L., Yan, R., Rao, H., Gao, G., Luo, S., 2014. Down-Regulation of Gli Transcription Factor Leads to the Inhibition of Migration and Invasion of Ovarian Cancer Cells via Integrin β4-Mediated FAK Signaling. PLoS ONE 9, e88386. https://doi.org/10.1371/journal.pone.0088386

Chiang, C., Litingtung, Y., Harris, M.P., Simandl, B.K., Li, Y., Beachy, P.A., Fallon, J.F., 2001. Manifestation of the Limb Prepattern: Limb Development in the Absence of Sonic Hedgehog Function. Developmental Biology 236, 421–435. https://doi.org/10.1006/dbio.2001.0346

Chiang, C., Litingtung, Y., Lee, E., Young, K.E., Corden, J.L., Westphal, H., Beachy, P.A., 1996. Cyclopia and defective axial patterning in mice lacking Sonic hedgehog gene function. Nature 383, 407–413. https://doi.org/10.1038/383407a0

Choi, K.-S., Lee, C., Harfe, B.D., 2012. Sonic hedgehog in the notochord is sufficient for patterning of the intervertebral discs. Mechanisms of development 129, 255–262. https://doi.org/10.1016/j.mod.2012.07.003

Dahn, R.D., Davis, M.C., Pappano, W.N., Shubin, N.H., 2006. Sonic hedgehog function in chondrichthyan fins and the evolution of appendage patterning. Nature 445, 311–314. https://doi.org/10.1038/nature05436

Dassule, H.R., Lewis, P., Bei, M., Maas, R., McMahon, A.P., 2000. Sonic hedgehog regulates growth and morphogenesis of the tooth. Development (Cambridge, England) 127, 4775–4785.

DeLaurier, A., Eames, B.F., BlanconSánchez, B., Peng, G., He, X., Swartz, M.E., Ullmann, B., Westerfield, M., Kimmel, C.B., 2010. Zebrafish sp7:EGFP: A transgenic for studying otic vesicle formation, skeletogenesis, and bone regeneration. genesis 48, 505–511. https://doi.org/10.1002/dvg.20639

Desvignes, T., Carey, A., Braasch, I., Enright, T., Postlethwait, J.H., 2018. Skeletal development in the heterocercal caudal fin of spotted gar (lepisosteus oculatus) and other lepisosteiformes. Developmental Dynamics 247, 724–740. https://doi.org/10.1002/dvdy.24617

Dworkin, S., Boglev, Y., Owens, H., Goldie, S.J., 2016. The Role of Sonic Hedgehog in Craniofacial Patterning, Morphogenesis and Cranial Neural Crest Survival. Journal of Developmental Biology 4, 24. https://doi.org/10.3390/jdb4030024

Ertzer, R., Müller, F., Hadzhiev, Y., Rathnam, S., Fischer, N., Rastegar, S., Strähle, U., 2007. Cooperation of sonic hedgehog enhancers in midline expression. Developmental Biology 301, 578–589. https://doi.org/10.1016/j.ydbio.2006.11.004

Ewald, A.J., Brenot, A., Duong, M., Chan, B.S., Werb, Z., 2008. Collective epithelial migration and cell rearrangements drive mammary branching morphogenesis. Developmental cell 14, 570–581. https://doi.org/10.1016/j.devcel.2008.03.003

Fernandes-Silva, H., Correia-Pinto, J., Moura, .R, 2017. Canonical Sonic Hedgehog Signaling in Early Lung Development. Journal of Developmental Biology 5, 3. https://doi.org/10.3390/jdb5010003

Fournier-Thibault, C., Blavet, C., Jarov, A., Bajanca, F., Thorsteinsdóttir, S., Duband, J.-L., 2009. Sonic Hedgehog Regulates Integrin Activity, Cadherin Contacts, and Cell Polarity to Orchestrate Neural Tube Morphogenesis. The Journal of Neuroscience 29, 12506–12520. https://doi.org/10.1523/jneurosci.2003-09.2009

Freitas, R., GómeznSkarmeta, J.L., Rodrigues, P.N., 2014. New frontiers in the evolution of fin development. Journal of Experimental Zoology Part B: Molecular and Developmental Evolution 322, 540–552. https://doi.org/10.1002/jez.b.22563

Freitas, R., Zhang, G., Cohn, M.J., 2006. Evidence that mechanisms of fin development evolved in the midline of early vertebrates. Nature 442, 1033–1037. https://doi.org/10.1038/nature04984

Goodrich, L. v, Johnson, R.L., Milenkovic, L., McMahon, J.A., Scott, M.P., 1996. Conservation of the hedgehog/patched signaling pathway from flies to mice: induction of a mouse patched gene by Hedgehog. Genes & Development 10, 301–312. https://doi.org/10.1101/gad.10.3.301

Hadzhiev, Y., Lele, Z., Schindler, S., Wilson, S.W., Ahlberg, P., Strähle, U., Müller, F., 2007. Hedgehog signaling patterns the outgrowth of unpaired skeletal appendages in zebrafish. BMC Developmental Biology 7, 75. https://doi.org/10.1186/1471-213X-7-75

Hanna, A., Shevde, L.A., 2016. Hedgehog signaling: modulation of cancer properies and tumor mircroenvironment. Molecular Cancer 15, 24. https://doi.org/10.1186/s12943-016-0509-3

Hu, D., Helms, J.A., 1999. The role of sonic hedgehog in normal and abnormal craniofacial morphogenesis. Development (Cambridge, England) 126, 4873–4884.

Hu, D., Young, N.M., Li, X., Xu, Y., Hallgrimsson, B., Marcucio, R.S., 2015. A dynamic Shh expression pattern, regulated by SHH and BMP signaling, coordinates fusion of primordia in the amniote face. Development 142, 567–574. https://doi.org/10.1242/dev.114835

Huang, P., Xiong, F., Megason, S.G., Schier, A.F., 2012. Attenuation of Notch and Hedgehog Signaling Is Required for Fate Specification in the Spinal Cord. PLoS Genetics 8, e1002762. https://doi.org/10.1371/journal.pgen.1002762

Huycke, T.R., Eames, B.F., Kimmel, C.B., 2012. Hedgehog-dependent proliferation drives modular growth during morphogenesis of a dermal bone. Development 139, 2371–2380. https://doi.org/10.1242/dev.079806

Ingham, P.W., 1993. Localized hedgehog activity controls spatial limits of wingless transcription in the Drosophila embryo. Nature 366, 560–562. https://doi.org/10.1038/366560a0

Jarov, A., Williams, K.P., Ling, L.E., Koteliansky, V.E., Duband, J.-L., Fournier-Thibault, C., 2003. A dual role for Sonic hedgehog in regulating adhesion and differentiation of neuroepithelial cells. Developmental Biology 261, 520–536. https://doi.org/10.1016/s0012-1606(03)00351-8 PMID - 14499657

Jaskoll, T., Leo, T., Witcher, D., Ormestad, M., Astorga, J., Bringas, P., Carlsson, P., Melnick, M., 2004. Sonic hedgehog signaling plays an essential role during embryonic salivary gland epithelial branching morphogenesis. Developmental Dynamics 229, 722–732. https://doi.org/10.1002/dvdy.10472

Jeong, J., Mao, J., Tenzen, T., Kottmann, A.H., McMahon, A.P., 2004. Hedgehog signaling in the neural crest cells regulates the patterning and growth of facial primordia. Genes & Development 18, 937–951. https://doi.org/10.1101/gad.1190304

Kim, H.Y., Pang, M.-F., Varner, V.D., Kojima, L., Miller, E., Radisky, D.C., Nelson, C.M., 2015. Localized Smooth Muscle Differentiation Is Essential for Epithelial Bifurcation during Branching Morphogenesis of the Mammalian Lung. Developmental Cell 34, 719– 726. https://doi.org/10.1016/j.devcel.2015.08.012

Krauss, R.S., Concordet, J., Ingham, P.W., 1993. A functionally conserved homolog of the Drosophila segment polarity gene hh is expressed in tissues with polarizing activity in zebrafish embryos. Cell 75, 1431–1444.

Laforest, L., Brown, C.W., Poleo, G., Géraudie, J., Tada, M., Ekker, M., Akimenko, M.A., 1998. Involvement of the sonic hedgehog, patched 1 and bmp2 genes in patterning of the zebrafish dermal fin rays. Development (Cambridge, England) 125, 4175–4184.

Larouche, O., Zelditch, M.L., Cloutier, .R, 2017. Fin modules: an evolutionary perspective on appendage disparity in basal vertebrates. BMC biology 15, 32. https://doi.org/10.1186/s12915-017-0370-x

Lee, H., Kimelman, D., 2002. A Dominant-Negative Form of p63 Is Required for Epidermal Proliferation in Zebrafish. Developmental Cell 2, 607–616.

Lee, Y., Hami, D., Val, S. de, Kagermeier-Schenk, B., Wills, A.A., Black, B.L., Weidinger, G., Poss, K.D., 2009. Maintenance of blastemal proliferation by functionally diverse epidermis in regenerating zebrafish fins. Developmental Biology 331, 270–280. https://doi.org/10.1016/j.ydbio.2009.05.545

Li, A., Cho, J.-H., Reid, B., Tseng, C.-C., He, L., Tan, P., Yeh, C.-Y., Wu, P., Li, Y., Widelitz, R.B., Zhou, Y., Zhao, M., Chow, R.H., Chuong, C.-M., 2018. Calcium oscillations coordinate feather mesenchymal cell movement by SHH dependent modulation of gap junction networks. Nature Communications 9, 5377. https://doi.org/10.1038/s41467-018-07661-5

Lin, T.L., Matsui, W., 2012. Hedgehog pathway as a drug target: Smoothened inhibitors in development. OncoTargets and therapy 5, 47–58. https://doi.org/10.2147/ott.s21957

Lorberbaum, D.S., Ramos, A.I., Peterson, K.A., Carpenter, B.S., Parker, D.S., De, S., Hillers, L.E., Blake, V.M., Nishi, Y., McFarlane, M.R., Chiang, A.C.Y., Kassis, J.A., Allen, B.L., McMahon, A.P., Barolo, S., 2016. An ancient yet flexible cis-regulatory architecture allows localized Hedgehog tuning by patched/Ptch1. eLife 5, e13550. https://doi.org/10.7554/elife.13550

Malik, S., 2012. Syndactyly: phenotypes, genetics and current classification. European Journal of Human Genetics 20, 817. https://doi.org/10.1038/ejhg.2012.14

Marigo, V., Scott, M.P., Johnson, R.L., Goodrich, L. v, Tabin, C.J., 1996. Conservation in hedgehog signaling: induction of a chicken patched homolog by Sonic hedgehog in the developing limb. Development (Cambridge, England) 122, 1225–1233.

Nakamura, T., Gehrke, A.R., Lemberg, J., Szymaszek, J., Shubin, N.H., 2016. Digits and fin rays share common developmental histories. Nature 537, 225. https://doi.org/10.1038/nature19322

Neumann, C.J., Grandel, H., Gaffield, W., Schulte-Merker, S., Nüsslein-Volhard, C., 1999. Transient establishment of anteroposterior polarity in the zebrafish pectoral fin bud in the absence of sonic hedgehog activity. Development (Cambridge, England) 126, 4817–4826.

Oster, G.F., Shubin, N., Murray, J.D., Alberch, P., 1988. EVOLUTION AND MORPHOGENETIC RULES: THE SHAPE OF THE VERTEBRATE LIMB IN

ONTOGENY AND PHYLOGENY. Evolution 42, 862–884. https://doi.org/10.1111/j.1558-5646.1988.tb02508.x

Pepicelli, C. v., Lewis, P.M., McMahon, A.P., 1998. Sonic hedgehog regulates branching morphogenesis in the mammalian lung. Current Biology 8, 1083–1086. https://doi.org/10.1016/S0960-9822(98)70446-4

Quint, E., Smith, A., Avaron, F., Laforest, L., Miles, J., Gaffield, W., Akimenko, M.-A., 2002. Bone patterning is altered in the regenerating zebrafish caudal fin after ectopic expression of sonic hedgehog and bmp2b or exposure to cyclopamine. Proceedings of the National Academy of Sciences 99, 8713–8718. https://doi.org/10.1073/pnas.122571799

Riccio, P., Cebrian, C., Zong, H., Hippenmeyer, S., Costantini, F., 2016. Ret and Etv4 Promote Directed Movements of Progenitor Cells during Renal Branching Morphogenesis. PLOS Biology 14, e1002382. https://doi.org/10.1371/journal.pbio.1002382

Riddle, R.D., Johnson, R.L., Laufer, E., Tabin, C.J., 1993. Sonic hedgehog mediates the polarizing activity of the ZPA. Cell 75, 1401–1416.

Rudolf, K., Umetsu, D., Aliee, M., Sui, L., Jülicher, F., Dahmann, C., 2015. A local difference in Hedgehog signal transduction increases mechanical cell bond tension and biases cell intercalations along the Drosophila anteroposterior compartment boundary. Development 142, 3845–3858. https://doi.org/10.1242/dev.125542

Sanders, T.A., Llagostera, E., Barna, M., 2013. Specialized filopodia direct long-range transport of SHH during vertebrate tissue patterning. Nature 497, 628. https://doi.org/10.1038/nature12157

Sato, N., Leopold, P.L., Crystal, R.G., 1999. Induction of the hair growth phase in postnatal mice by localized transient expression of Sonic hedgehog. Journal of Clinical Investigation 104, 855–864. https://doi.org/10.1172/jci7691

Saunders, J.W., Gasseling, M.T., 1968. Ectoderm-mesenchymal interactions in the origin of wing symmetry. Epithelial-Mesenchymal Interactions 78–97.

Sehring, I.M., Weidinger, G., 2019. Recent advancements in understanding fin regeneration in zebrafish. WIREs Developmental Biology 9, e367. https://doi.org/10.1002/wdev.367

Seppala, M., Fraser, G.J., Birjandi, A.A., Xavier, G.M., Cobourne, M.T., 2017. Sonic Hedgehog Signaling and Development of the Dentition. Journal of Developmental Biology 5, 6. https://doi.org/10.3390/jdb5020006

Shibata, E., Ando, K., Murase, E., Kawakami, A., 2018. Heterogeneous fates and dynamic rearrangement of regenerative epidermis-derived cells during zebrafish fin regeneration. Development 145, dev.162016. https://doi.org/10.1242/dev.162016

Shkumatava, A., 2004. Sonic hedgehog, secreted by amacrine cells, acts as a short-range signal to direct differentiation and lamination in the zebrafish retina. Development 131, 3849– 3858. https://doi.org/10.1242/dev.01247

Shubin, N.H., Alberch, P., 1986. A morphogenetic approach to the origin and basic organization of the tetrapod limb. Evolutionary Biology 319–387. https://doi.org/10.1007/978-1-4615-6983-1_6

Spurlin, J.W., Nelson, C.M., 2017. Building branched tissue structures: from single cell guidance to coordinated construction. Philosophical Transactions of the Royal Society B: Biological Sciences 372, 20150527. https://doi.org/10.1098/rstb.2015.0527

Stewart, S., Gomez, A.W., Armstrong, B.E., Henner, A., Stankunas, K., 2014. Sequential and Opposing Activities of Wnt and BMP Coordinate Zebrafish Bone Regeneration. Cell reports 6, 482–498. https://doi.org/10.1016/j.celrep.2014.01.010

Swartz, M.E., Nguyen, V., McCarthy, N.Q., Eberhart, J.K., 2012. Hh signaling regulates patterning and morphogenesis of the pharyngeal arch-derived skeleton. Developmental biology 369, 65–75. https://doi.org/10.1016/j.ydbio.2012.05.032

Testaz, S., Jarov, A., Williams, K.P., Ling, L.E., Koteliansky, V.E., Fournier-Thibault, C., Duband, J.-L., 2001. Sonic hedgehog restricts adhesion and migration of neural crest cells independently of the Patched-Smoothened-Gli signaling pathway. Proceedings of the National Academy of Sciences 98, 12521–12526. https://doi.org/10.1073/pnas.221108698

Tickle, C., Towers, M., 2017. Sonic Hedgehog Signaling in Limb Development. Frontiers in Cell and Developmental Biology 5, 14. https://doi.org/10.3389/fcell.2017.00014

Tsai, T.Y.-C., Sikora, M., Xia, P., Colak-Champollion, T., Knaut, H., Heisenberg, C.-P., Megason, S.G., 2020. An adhesion code ensures robust pattern formation during tissue morphogenesis. Science 370, 113–116. https://doi.org/10.1126/science.aba6637

Woo, W.-M., Zhen, H.H., Oro, A.E., 2012. Shh maintains dermal papilla identity and hair morphogenesis via a Noggin–Shh regulatory loop. Genes & Development 26, 1235–1246. https://doi.org/10.1101/gad.187401.112

Zeng, C., Chen, T., Zhang, Y., Chen, Q., 2017. Hedgehog signaling pathway regulates ovarian cancer invasion and migration via adhesion molecule CD24. Journal of Cancer 8, 786–792. https://doi.org/10.7150/jca.17712

Zhang, J., Jeradi, S., Strähle, U., Akimenko, M.-A., 2012b. Laser ablation of the sonic hedgehog-a-expressing cells during fin regeneration affects ray branching morphogenesis. Developmental Biology 365, 424–433. https://doi.org/10.1016/j.ydbio.2012.03.008

Zuniga, A., 2015. Next generation limb development and evolution: old questions, new perspectives. Development 142, 3810–3820. https://doi.org/10.1242/dev.125757

